# A novel heteromeric pantothenate kinase complex in apicomplexan parasites

**DOI:** 10.1101/2021.03.16.435557

**Authors:** Erick T. Tjhin, Vanessa M. Howieson, Christina Spry, Giel G. van Dooren, Kevin J. Saliba

## Abstract

Coenzyme A is synthesised from pantothenate via five enzyme-mediated steps. The first step is catalysed by pantothenate kinase (PanK). All PanKs characterised to date form homodimers. Many organisms express multiple PanKs. In some cases, these PanKs are not functionally redundant, and some appear to be non-functional. Here, we investigate the PanKs in two pathogenic apicomplexan parasites, *Plasmodium falciparum* and *Toxoplasma gondii*. Each of these organisms express two PanK homologues (PanK1 and PanK2). We demonstrate that *Pf*PanK1 and *Pf*PanK2 associate, forming a single, functional PanK complex that includes the multi-functional protein, *Pf*14-3-3I. Similarly, we demonstrate that *Tg*PanK1 and *Tg*PanK2 form a single complex that possesses PanK activity. Both *Tg*PanK1 and *Tg*PanK2 are essential for *T. gondii* proliferation, specifically due to their PanK activity. Our study constitutes the first examples of heteromeric PanK complexes in nature and provides an explanation for the presence of multiple PanKs within certain organisms.

## INTRODUCTION

Coenzyme A (CoA) is an essential enzyme cofactor in all living organisms ^1^. CoA itself, and the CoA-derived phosphopantetheine prosthetic group required by various carrier proteins, function as acyl group carriers and activators in key cellular processes such as fatty acid biosynthesis, β-oxidation and the citric acid cycle. Pantothenate kinase (PanK) catalyses the first step in the conversion of pantothenate (vitamin B5) to CoA ^2^. PanKs are categorised into three distinct types, type I, II and III based on their primary sequences, structural fold, enzyme kinetics and inhibitor sensitivity. PanKs from all three types have been shown to exist as homodimers based on their solved protein structures ^3–10^. All eukaryotic PanKs that have been characterised so far are type II PanKs ^5^. Interestingly, many eukaryotes express multiple PanKs (such as *Arabidopsis thaliana* ^11, 12^, *Mus musculus* ^13–16^ and *Homo sapiens* ^17–21^), and in some cases it is clear that these PanKs are not functionally redundant ^15, 22^. For example, mutations in only one of four type II PanKs in humans causes a neurodegenerative disorder known as PanK-associated neurodegeneration ^17^. Some bacteria also express multiple PanKs. For example, some *Mycobacterium* ^23^, *Streptomyces* ^7^ and *Bacillus* ^7, 24, 25^ species have both type I and type III PanKs, while a select few bacilli (including the category A biodefense pathogen *Bacillus anthracis*) carry both a type II and type III PanK ^7^. In some organisms harbouring multiple PanKs, it has not been possible to demonstrate functional activity for all enzymes. One of the four human type II PanKs is shown to be catalytically inactive^21 26^, as is a type III PanK from *Mycobacterium tuberculosis* ^23^, and a type II PanK from *B. anthracis* ^7^. The reason for the presence of multiple PanKs within certain cells, and the apparent inactivity of certain PanKs, is unclear.

Two putative genes coding for PanK enzymes have been identified in each of the genomes of the pathogenic apicomplexan parasites *Plasmodium falciparum* (PF3D7_1420600 (*Pfpank1*) and PF3D7_1437400 (*Pfpank2*)) and *Toxoplasma gondii* (TGME49_307770 (*Tgpank1*) and TGME49_235478 (*Tgpank2*)). We have recently shown that mutations in *Pf*PanK1 alter PanK activity in *P. falciparum*, providing evidence that *Pf*PanK1 is an active PanK, at least in the disease-causing stage of the parasite’s lifecycle ^27^. The function of *Pf*PanK2, and its contribution to PanK activity in *P. falciparum*, is unknown. *Pf*PanK2 contains a unique, large insert in a loop associated with the dimerisation of PanKs in their native conformation ^8^ and this may affect its ability to form a dimer, rendering it inactive ^28^. No functional information is available on the putative *T. gondii* PanKs, but a genome-wide CRISPR-Cas9 screen of the *T. gondii* genome predicted that *both* PanK genes are important for parasite growth *in vitro* ^29^. Similarly, a recent genome-wide insertional mutagenesis study of *P. falciparum* has predicted *both Pf*PanK1 and *Pf*PanK2 to be essential ^30^. These results suggest that the PanK2 proteins of these parasites play important role(s), although their exact function remains unclear.

In this study, we demonstrate that PanK1 and PanK2 from *P. falciparum* and *T. gondii* are part of the same, multimeric protein complex. This constitutes the first identification of a heteromeric PanK complex in nature. Furthermore, our data provide the first evidence that PanK2 is essential for PanK function in apicomplexans.

## RESULTS

### *Pf*PanK1 and *Pf*PanK2 are part of the same protein complex

The importance and role of *Pf*PanK2 in apicomplexan parasites have not previously been established. To characterise the *P. falciparum* PanK2 homologue (*Pf*PanK2), we first determined where in the parasite the protein localises. We episomally expressed *Pf*PanK2-GFP in asexual blood stage *P. falciparum* parasites and found that *Pf*PanK2-GFP is localised throughout the parasite cytosol and is not excluded from the nucleus (**Figure 1a**). This is a similar localisation to what we observed for *Pf*PanK1-GFP previously ^27^. Western blotting of proteins separated by SDS-PAGE revealed that *Pf*PanK2-GFP has a molecular mass consistent with the predicted mass of the fusion protein (∼118 kDa; **Figure 1b**), which is slightly higher than the predicted mass of *Pf*PanK1-GFP (∼87 kDa; **Figure 1b**). As PanKs from other organisms exist as homodimers, we undertook blue native-PAGE to determine whether *Pf*PanK1-GFP and *Pf*PanK2-GFP exist in protein complexes. Interestingly, under native conditions, both *Pf*PanK1-GFP and *Pf*PanK2-GFP were found to be part of complexes that are ∼240 kDa in mass (**Figure 1b**).

**Figure 1.**
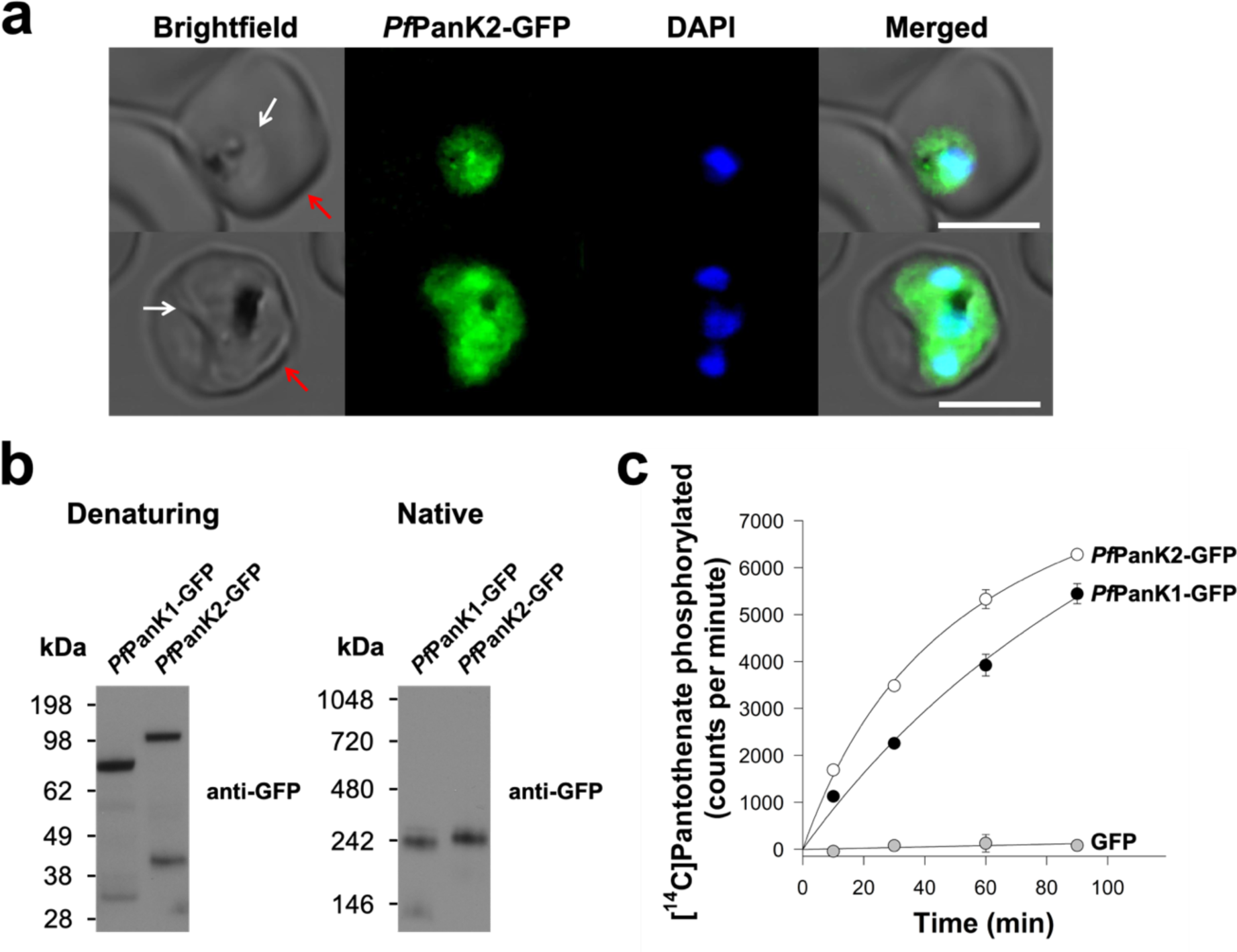
*Pf*PanK1 and *Pf*PanK2 are part of similar-sized protein complexes that possess PanK activity. (a) Confocal micrographs showing the subcellular location of *Pf*PanK2-GFP within trophozoite/schizont-stage *P. falciparum*-infected erythrocytes. The nuclei of the parasites are labelled with DAPI. From left to right: Brightfield, GFP-fluorescence, DAPI-fluorescence, and merged images. Arrows indicate the plasma membranes of the erythrocyte (red) or the parasite (white). Scale bars represent 5 µm. (b) Denaturing and native western blot analyses of the GFP-tagged proteins in *Pf*PanK1-GFP and *Pf*PanK2-GFP expressing parasites. The expected sizes of the proteins are ∼87 kDa for *Pf*PanK1-GFP and ∼118 kDa for *Pf*PanK2-GFP. The molecular mass of the GFP tag is ∼27 kDa. Western blots were performed with anti-GFP antibodies and each of the blots shown is representative of three independent experiments, each performed with a different batch of parasites. (c) The phosphorylation of [^14^C]pantothenate (initial concentration of 2 µM, ∼10,000 counts per minute) over time by the immunopurified complex from lysates of parasites expressing *Pf*PanK1-GFP (black circles), *Pf*PanK2-GFP (white circles) and untagged GFP (grey circles). Data shown are representative of two independent experiments, each performed with a different batch of parasites and carried out in duplicate. Error bars represent range/2 and are not shown if smaller than the symbols.

To determine the activity and protein composition of these complexes, we set out to purify the *Pf*PanK1-GFP and *Pf*PanK2-GFP complexes by immunoprecipitation. As a control, we also purified untagged GFP. We verified that most of the GFP-tagged proteins were captured from the total lysates prepared from the different cell lines, with bands corresponding to *Pf*PanK1-GFP, *Pf*PanK2-GFP and the untagged GFP epitope tag detected in the bound fraction of the respective cell lines (**Figure S1**). To determine whether the purified *Pf*PanK1 and *Pf*PanK2 complexes possess PanK activity, we performed a [^14^C]pantothenate phosphorylation assay. We found that 50 − 60% of the [^14^C]pantothenate initially present in the reaction was phosphorylated within 90 min by the immunopurified complex from *both* the *Pf*PanK1-GFP- and *Pf*PanK2-GFP-expressing lines (**Figure 1c**). Conversely, the immunopurified untagged GFP did not display PanK activity (**Figure 1c**). These experiments provide the first indication that *Pf*PanK1 and *Pf*PanK2 are part of an active PanK enzyme complex in *P. falciparum* parasites. They also provide the first indication that *Pf*PanK2 contributes to PanK activity in these parasites.

To elucidate the protein composition of the *Pf*PanK1-GFP and *Pf*PanK2-GFP complexes, the immunoprecipitated samples were subjected to mass spectrometry (MS)-based proteomic analysis (bound fractions of untagged GFP-expressing and 3D7 wild-type parasites were included as negative controls). Both *Pf*PanK1 (36 – 50% coverage, **Figure S2**) and *Pf*PanK2 (29 – 49% coverage, **Figure S3**) were unequivocally detected as the two most abundant proteins in the immunopurified complex from *both* the *Pf*PanK1-GFP- and *Pf*PanK2-GFP-expressing cells (**Figure 2a**). Interestingly the next most abundant protein detected in both complexes was *Pf*14-3-3I (43 – 67% coverage, **Figure 2a** and **Figure S4**). These results are consistent with *Pf*PanK1, *Pf*PanK2 and *Pf*14-3-3I being part of the same protein complex. Other proteins, such as M17 leucyl aminopeptidase (fourth most abundant), were also detected in the MS analysis, albeit with a comparatively fewer number of peptides (**Figure 2a**, and **Table S3**).

**Figure 2.**
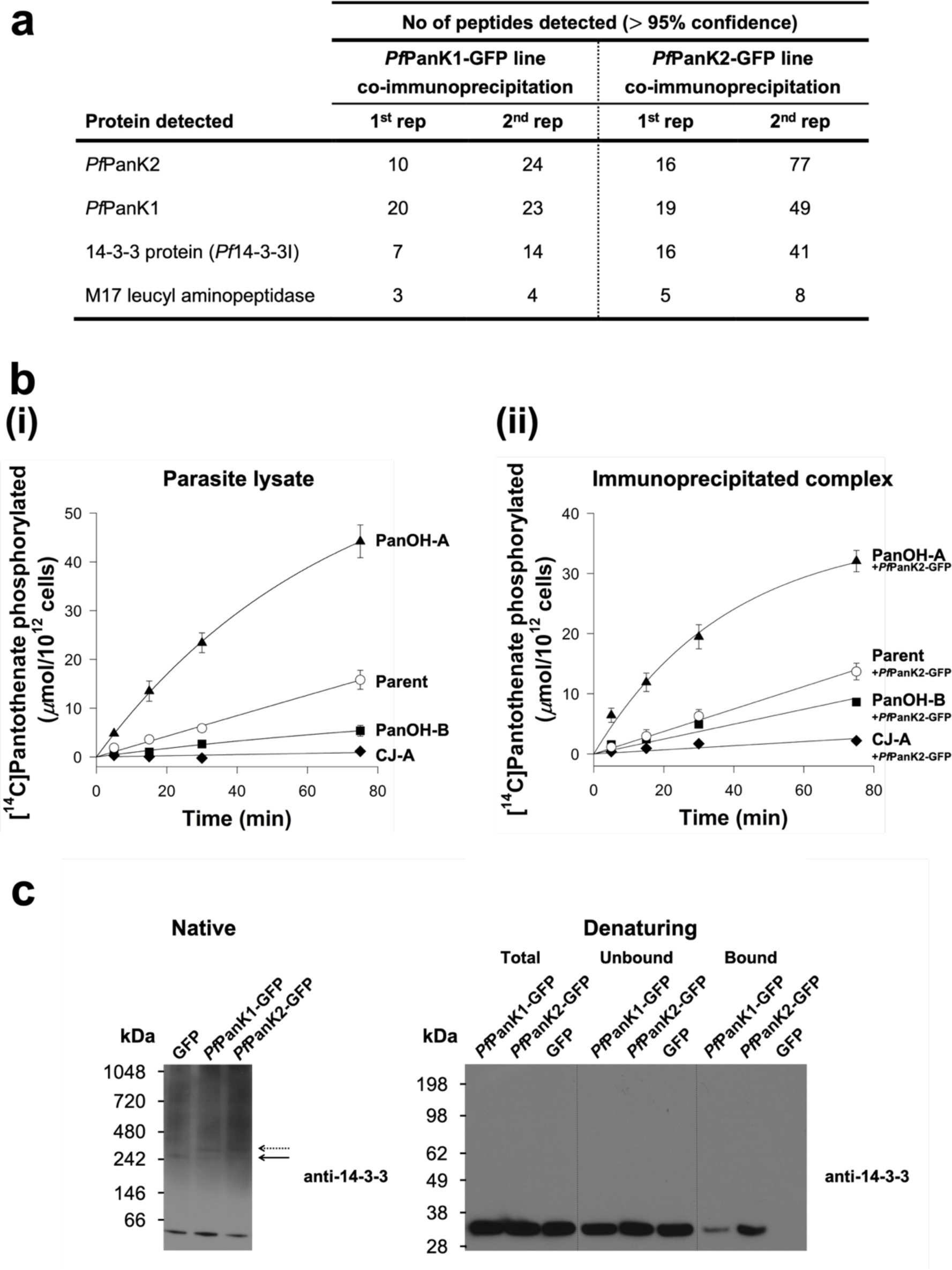
*Pf*PanK1 and *Pf*PanK2 are part of a single PanK complex that includes *Pf*14-3-3I. (a) The four most abundant proteins identified in the MS analysis of proteins immunoprecipitated with anti-GFP antibodies from *Pf*PanK1-GFP- and *Pf*PanK2-GFP-expressing parasites. Data shown are representative of two independent analyses (1^st^ and 2^nd^ rep), each performed with a different batch of parasites. Only proteins with three or more peptides detected in both replicate co-immunoprecipitation experiments are shown. Proteins detected in the untagged GFP-expressing or wild-type 3D7 parasite immunoprecipitations (negative controls) were removed. Proteins are listed in descending order according to the total number of peptides detected across all replicates (all four column). (b) The phosphorylation of [^14^C]pantothenate (initial concentration of 2 µM) over time by (i) lysates generated from Parent (white circles), PanOH-A (black triangles), PanOH-B (black squares) and CJ-A (black diamonds) parasites (reproduced from ^27^) and (ii) proteins immunoprecipitated with anti-GFP antibodies from Parent^+*Pf*PanK2-GFP^ (white circles), PanOH-A^+*Pf*PanK2-GFP^ (black triangles), PanOH-B^+*Pf*PanK2-GFP^ (black squares) and CJ-A^+*Pf*PanK2-GFP^ (black diamonds) parasite lysates. Values in (ii) are averaged from three independent experiments, each performed with a different batch of parasites and carried out in duplicate. Error bars represent SEM and are not shown if smaller than the symbols. (c) Native western blot analysis of the lysates and denaturing western blot analyses of the different GFP-trap co-immunoprecipitation fractions of *Pf*PanK1-GFP- and *Pf*PanK2-GFP-expressing parasites, with untagged GFP-expressing parasite as a control. Western blots were performed with pan-specific anti-14-3-3 antibodies (previously shown to detect *Plasmodium* 14-3-3 ^32^). Arrows indicate the position of 14-3-3-containing complexes of comparable masses to the complexes found in *Pf*PanK1-GFP- and *Pf*PanK2-GFP-expressing parasites. The native blot shown is a representative of three independent experiments, while the denaturing blot is a representative of two independent experiments, each performed with a different batch of parasites.

To test further whether *Pf*PanK1 and *Pf*PanK2 are part of the same protein complex, we introduced episomally-expressed *Pf*PanK2-GFP into parasite strains generated in a previous study ^27^. These mutant strains, termed PanOH-A, PanOH-B and CJ-A, were generated by drug-pressuring parasites with antiplasmodial pantothenate analogues, and harbour mutations in *Pf*PanK1 that affect *Pf*PanK catalytic activity ^27^. We immunopurified the *Pf*PanK2-GFP complex from the PanOH-A, PanOH-B and CJ-A strains, as well as from wild type (Parent) parasites that expresses *Pf*PanK2-GFP as a control, and performed [^14^C]pantothenate phosphorylation assays on the immunopurified *Pf*PanK2-GFP complex. As we reported previously, the *Pf*PanK1 mutations alter the *Pf*PanK activity of PanOH-A, PanOH-B and CJ-A parasites such that the following rank order of enzyme activity relative to the Parent line is observed: PanOH-A > Parent > PanOH-B > CJ-A (**Figure 2b(i),** ^27^). Notably, PanK activity of the immunopurified *Pf*PanK2-GFP complex from the various *Pf*PanK2-GFP expressing lines followed the same rank order (i.e. PanOH-A^+*Pf*PanK2-GFP^ > Parent^+*Pf*PanK2-GFP^ > PanOH-B^+*Pf*PanK2-GFP^ > CJ-A^+*Pf*PanK2-GFP^). This difference in pantothenate phosphorylation rates was not due to variations in the amount of *Pf*PanK2-GFP protein in the immunopurified complexes used for the assays (**Figure S5**). These data are consistent with *Pf*PanK2-GFP associating with the mutant *Pf*PanK1 from each cell line and indicate that both proteins are part of the same PanK complex in *Plasmodium* parasites.

Our proteomic analysis identified *Pf*14-3-3I as being co-immunoprecipitated with both *Pf*PanK1 and *Pf*PanK2 (**Figure 2a**). To test whether *Pf*14-3-3I is a *bona fide* component of the PanK complex of *P. falciparum,* we performed western blotting with a pan-specific anti-14-3-3 antibody. Under native conditions, the 14-3-3 antibody detected a major protein band at <66 kDa, which likely represents dimeric *Pf*14-3-3I proteins from the parasite ^31^. We also observed a protein complex of ∼240 kDa in each of the *Pf*PanK1-GFP-, *Pf*PanK2-GFP- and untagged GFP-expressing parasites (solid arrow, **Figure 2c**). In addition, a protein complex of slightly higher molecular mass, likely corresponding to the PanK complex that includes the GFP epitope tag, was also observed in the *Pf*PanK1-GFP-, *Pf*PanK2-GFP-expressing parasites but not the untagged GFP-expressing parasites (dashed arrow, **Figure 2c**). As a direct test for whether *Pf*14-3-3I exists in the same complex as *Pf*PanK1 and *Pf*PanK2, we performed western blotting on proteins immunopurified with anti-GFP antibodies from the *Pf*PanK1-GFP-, *Pf*PanK2-GFP- and untagged GFP-expressing parasites. We found that *Pf*14-3-3I protein was detected in the immunopurified complex from both the *Pf*PanK1-GFP- and *Pf*PanK2-GFP-expressing cells, but not in that purified from untagged GFP-expressing cells (**Figure 2c**). Together with the native western blot (**Figure 1b**) and proteomic (**Figure 2a**) analyses, these results are consistent with *Pf*PanK1 and *Pf*PanK2 being part of the *same* complex that also contains *Pf*14-3-3I, and that this complex is responsible for the PanK activity observed in the intraerythrocytic stage of *P. falciparum*.

### *Tg*PanK1 and *Tg*PanK2 also constitute a single complex with PanK activity that is essential for parasite proliferation

Based on sequence similarity, *Tg*PanK1 and *Tg*PanK2 are homologous to their *P. falciparum* counterparts (**Figure S6**). To begin to characterise *Tg*PanK1 and *Tg*PanK2, we introduced the coding sequence for a mini-Auxin-Inducible Degron (mAID)-haemagglutinin (HA) tag into the 3’ region of the open reading frames of *Tg*PanK1 or *Tg*PanK2 in RH Δ*Ku80*:TIR1 strain *T. gondii* parasites ^33^ also expressing a ‘tdTomato’ red fluorescent protein (RFP). Gene models with inserts sizes are noted (**Figure S7a**), and successful integration of the mAIDHA tag was verified by PCR (**Figure S7b**). Western blotting revealed that the *Tg*PanK1-mAIDHA and *Tg*PanK2-mAIDHA proteins have molecular masses of ∼160 and ∼200 kDa, respectively (**Figure 3a**), corresponding to the predicted sizes of *Tg*PanK1-mAIDHA (141 kDa) and *Tg*PanK2-mAIDHA (187 kDa). When analysed under native conditions, *Tg*PanK1-mAIDHA and *Tg*PanK2-mAIDHA both exist in protein complexes of ∼720 kDa in mass (**Figure 3a**).

**Figure 3.**
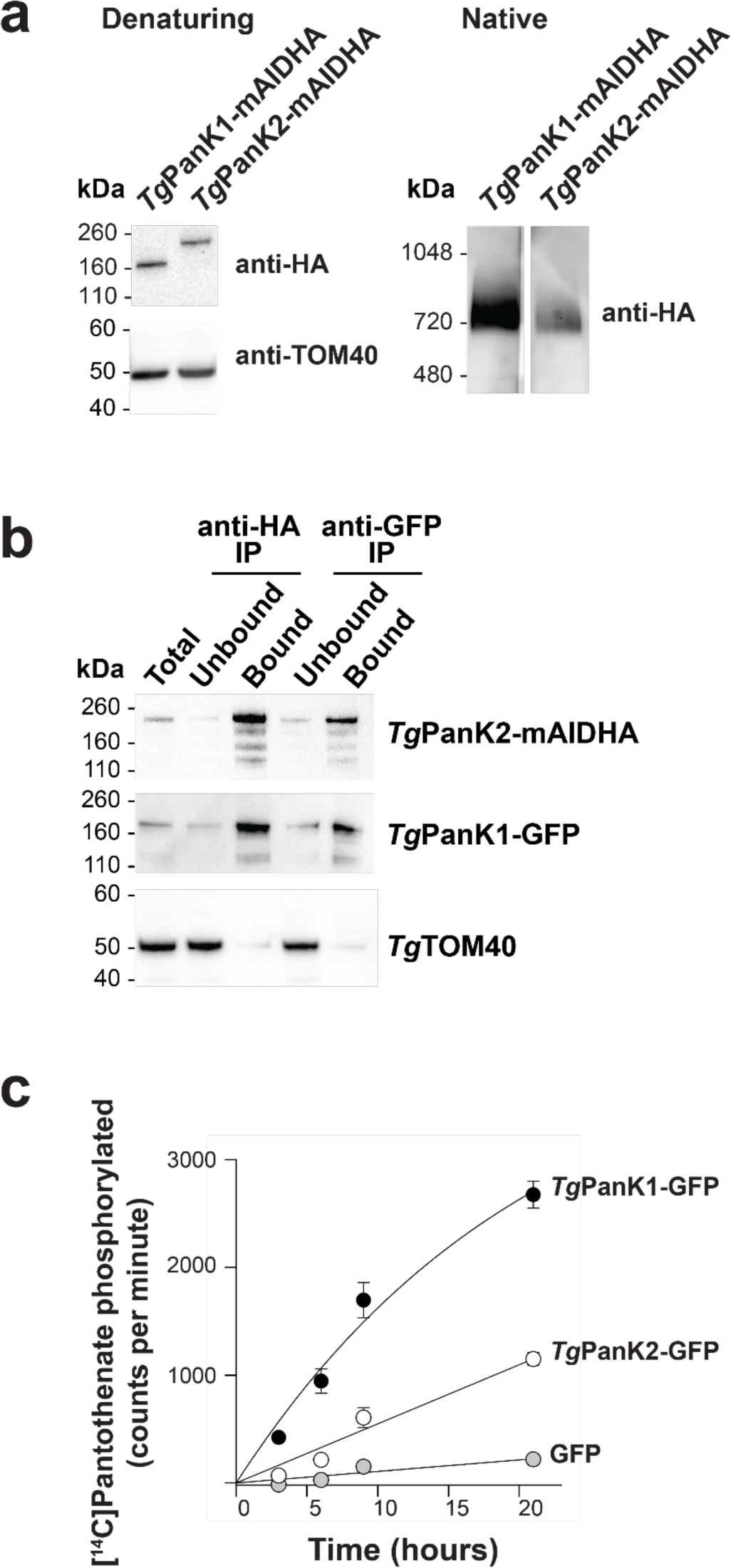
*Tg*PanK1 and *Tg*PanK2 are part of a single protein complex with PanK activity. (a) Denaturing and native western blot analyses of the HA-tagged proteins in *Tg*PanK1-mAIDHA and *Tg*PanK2-mAIDHA expressing parasites. The expected sizes of *Tg*PanK1-mAIDHA and *Tg*PanK2-mAIDHA are ∼141 kDa and ∼187 kDa, respectively. Western blots were performed with an anti-HA antibody and each blot is representative of three independent experiments, each performed with different batches of parasites. Denaturing western blots were also probed with anti-*Tg*TOM40, which served as a loading control. (b) Western blot analysis of proteins from *Tg*PanK1-GFP/*Tg*PanK2-mAIDHA parasite lysates immunoprecipitated with GFP-Trap and anti-HA beads. Protein samples were collected before immunoprecipitation (Total), from the fraction not bound to the GFP-Trap nor anti-HA beads (Unbound), and from the fraction bound to the GFP-Trap/anti-HA beads (Bound). Membranes were probed with anti-GFP and anti-HA antibodies, and the blot shown is representative of three independent experiments, each performed with different batches of parasites. *Tg*TOM40 served as a control protein that is part of an unrelated protein complex. Bound fractions contain protein from 4 × as many cells as the total and unbound lanes. (c) The phosphorylation of [^14^C]pantothenate (initial concentration 2 µM) over time by protein samples immunoprecipitated with GFP-Trap from parasites expressing *Tg*PanK1-GFP/*Tg*PanK2-mAIDHA (black circles),*Tg*PanK1-HA/*Tg*PanK2-GFP (white circles) and untagged GFP (grey circles). Data shown are representative of two independent experiments, each performed with a different batch of parasites and carried out in duplicate. Error bars represent range/2 and are not shown if smaller than the symbols.

To investigate if *Tg*PanK1 and *Tg*PanK2 are part of the same ∼720 kDa complex, we introduced a sequence encoding a GFP tag into the genomic locus of *Tg*PanK1 in the *Tg*PanK2-mAIDHA strain (integration verified by PCR (**Figure S7a** and **S7c**)). Co-immunoprecipitation experiments revealed that *Tg*PanK1-GFP co-purified with *Tg*PanK2-mAIDHA (**Figure 3b**). Analogous experiments with a *Tg*PanK1-HA/*Tg*PanK2-GFP line, wherein we integrated a sequence encoding a GFP tag into the *Tg*PanK2 locus and a sequence encoding a HA tag into the *Tg*PanK1 locus (integration verified by PCR (**Figure S7a** and **S7d**)), yielded similar results **(Figure S8)**. We therefore conclude that, like *Pf*PanK1 and *Pf*PanK2 in *P. falciparum* (**Figures 1 and 2**), *Tg*PanK1 and *Tg*PanK2 are components of the same protein complex.

To determine whether the *Tg*PanK1/*Tg*PanK2 complex has pantothenate kinase activity, we immunopurified proteins from *Tg*PanK1-GFP/*Tg*PanK2-mAIDHA, *Tg*PanK1-HA/*Tg*PanK2-GFP and control (expressing untagged GFP) cell lines using GFP-Trap, and measured the ability of the purified proteins to phosphorylate pantothenate. The samples purified from the *Tg*PanK1-GFP/*Tg*PanK2-mAIDHA and *Tg*PanK1-HA/*Tg*PanK2-GFP lines exhibited higher pantothenate phosphorylation activity than that from the control untagged GFP-expressing line (**Figure 3c**). These findings indicate that, like the *P. falciparum* PanKs (**Figure 1 and 2**), the *Tg*PanK1/ *Tg*PanK2 complex possesses PanK activity.

As is the case for *Pf*PanK2, the nucleotide-binding motifs of *Tg*PanK2 deviate substantially from those of other eukaryotic PanKs (**Figure S6**). It is therefore unclear whether pantothenate phosphorylation is catalysed solely by *Tg*PanK1 or if *Tg*PanK2 also contributes to PanK activity. To answer this, we first investigated whether *Tg*PanK1 and *Tg*PanK2 are important for parasite proliferation. *Tg*PanK1 and *Tg*PanK2 were individually knocked down by exposing the mAID-regulated lines to 100 µM indole-3-acetic acid (IAA – a plant hormone of the auxin class), a concentration that we determined was not detrimental to wild-type parasite proliferation. *Tg*PanK1-mAIDHA and *Tg*PanK2-mAIDHA were degraded within an hour of exposing the parasites to IAA (**Figure 4a**). Both the *Tg*PanK1-mAIDHA and *Tg*PanK2-mAIDHA lines express RFP, which enabled us to monitor parasite proliferation using fluorescence growth assays, as described previously ^34^. We measured proliferation of the *Tg*PanK1-mAIDHA, *Tg*PanK2-mAIDHA and parental lines cultured in the presence or absence of 100 µM IAA over seven days. In the absence of IAA, we observed a normal sigmoidal growth curve for all three strains (**Figure 4b**). By contrast, we observed a complete cessation of proliferation of both the *Tg*PanK1-mAIDHA- and *Tg*PanK2-mAIDHA-expressing parasite lines, but not the parental strain, in the presence of 100 µM IAA (**Figure 4b**). These data indicate that both *Tg*PanK1 and *Tg*PanK2 are crucial for *T. gondii* proliferation and, notably, that neither can substitute for the other. To establish whether *Tg*PanK1 and *Tg*PanK2 are essential due to the PanK activity of the complex, we constitutively expressed the type II PanK of *Staphylococcus aureus* (*Sapank)* in both the *Tg*PanK1-mAIDHA- and *Tg*PanK2-mAIDHA-expressing parasite lines, generating lines that we termed *Tg*PanK1-mAIDHA^+*Sa*PanK-Ty1^ and *Tg*PanK2-mAIDHA^+*Sa*PanK-Ty1^. The expression of the *Sa*PanK protein in these strains was verified by immunofluorescence microscopy and western blot (**Figure S9**). We measured the proliferation of the *Tg*PanK1-mAIDHA^+*Sa*PanK-Ty1^ and *Tg*PanK2-mAIDHA^+*Sa*PanK-Ty1^ lines in the presence and absence of 100 µM IAA, and compared this with expression of the *Tg*PanK1-mAIDHA, *Tg*PanK2-mAIDHA and parental lines. We obtained fluorescence measurements over a 7 day period, and compared the proliferation of each strain when the parental strain cultured in the absence of IAA reached mid-log phase. We found that both the *Tg*PanK1-mAIDHA^+*Sa*PanK-Ty1^ and *Tg*PanK2-mAIDHA^+*Sa*PanK-Ty1^ lines proliferated at a similar rate to the parental control line when cultured in the presence of IAA, in contrast to the *Tg*PanK1-mAIDHA and *Tg*PanK2-mAIDHA lines, where minimal proliferation was observed (**Figure 4c**). Collectively, our studies on *Tg*PanK1 and *Tg*PanK2 reveal that (i) *Tg*PanK1 and *Tg*PanK2 are part of the same protein complex, (ii) expression of both is required for PanK activity, and (iii) PanK activity of the complex is important for *T. gondii* proliferation during the disease-causing tachyzoite stage.

**Figure 4.**
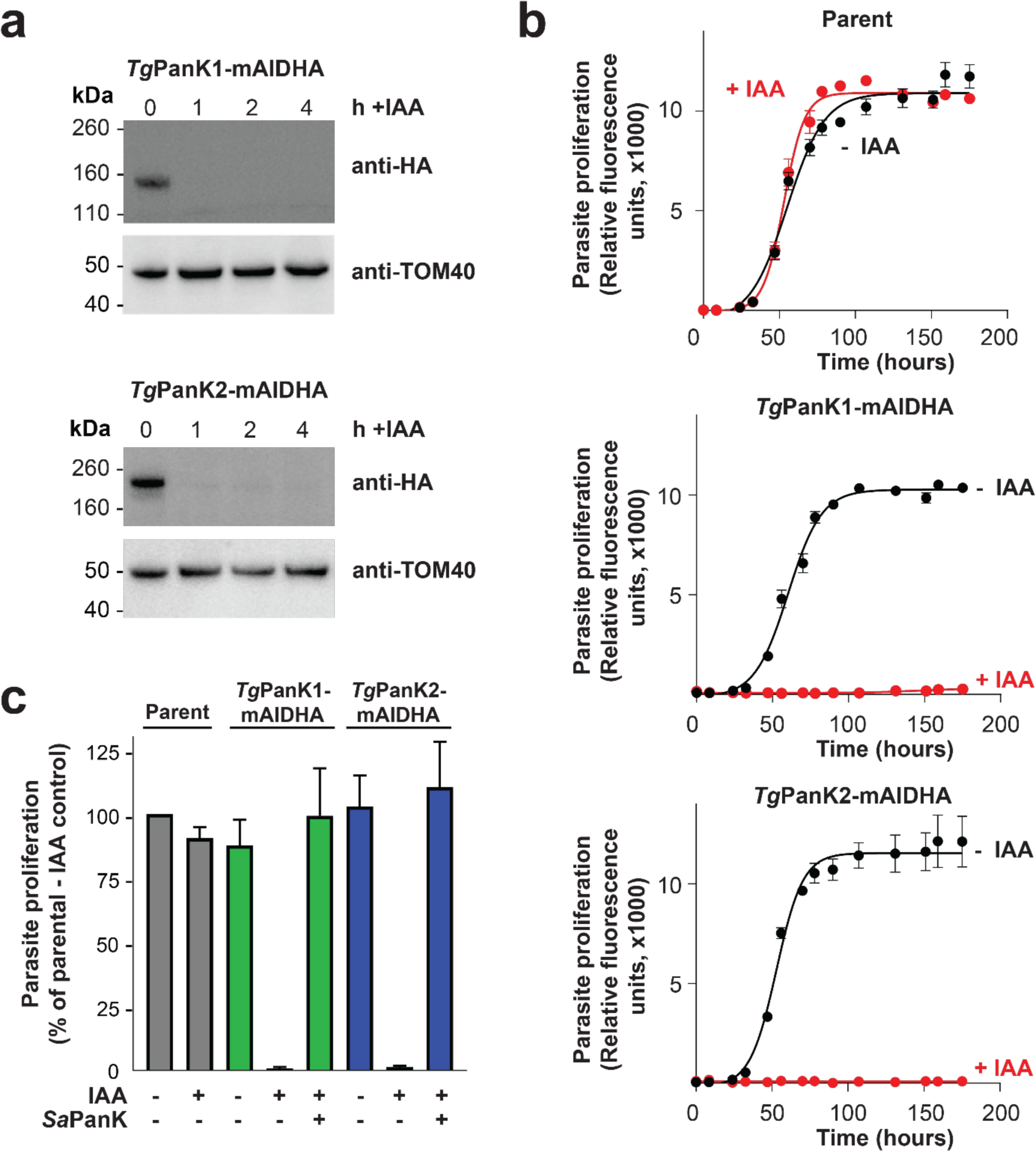
Expression of both *Tg*PanK1 and *Tg*PanK2 is necessary for PanK activity and for *T. gondii* tachyzoite proliferation. (a) IAA-induced knockdown of *Tg*PanK1-mAIDHA or *Tg*PanK2-mAIDHA protein over time. Western blot analysis of *Tg*PanK1-mAIDHA- and *Tg*PanK2-mAIDHA-expressing parasites, either in the absence of, or after 1, 2 and 4 hours of exposure to 100 µM IAA. Membranes were probed with anti-HA antibody to detect the *Tg*PanK1-mAIDHA and *Tg*PanK2-mAIDHA proteins, and anti-*Tg*Tom40 as a loading control. Western blots are representative of three independent experiments, each performed with a different batch of parasites. (b) The effect of *Tg*PanK1-mAIDHA or *Tg*PanK2-mAIDHA knockdown on *T. gondii* tachyzoite proliferation. tdTomato RFP-expressing parasites (Parent, *Tg*PanK1-mAIDHA and *Tg*PanK2-mAIDHA) were cultured over 7 days in the presence (red circles) or absence (black circles) of 100 µM IAA. Parasite proliferation was measured over time by assessing the RFP expression using a fluorescence reader. Graphs shown are representative of three independent experiments carried out in triplicate, each performed with a different batch of parasites. Error bars represent SD and are not shown if smaller than the symbols. (c) Complementation of *Tg*PanK1 and *Tg*PanK2 knockdown with *S. aureus* PanK. *S. aureus* PanK was constitutively expressed in *Tg*PanK1-mAIDHA (green bars) and *Tg*PanK2-mAIDHA (blue bars) parasites. The Parent RH RFP (grey bars), non-complemented and *Sa*PanK-Ty1-complemented parasites were cultured alongside the *Tg*PanK1-mAIDHA and *Tg*PanK2-mAIDHA lines in the presence (+) or absence (-) of 100 µM IAA. Parasite proliferation was monitored 1-2 times daily for 7 days. Proliferation was compared when the parental strain cultured in the absence of IAA was at the mid-log phase of parasite proliferation. Values are averaged from three independent experiments, each performed with a different batch of parasites and carried out in triplicate. Error bars represent SEM.

## DISCUSSION

All PanKs characterised to date have been shown to exist as homodimers ^3–10^. Here we present data consistent with PanK1 and PanK2 of the apicomplexan parasites *P. falciparum* and *T. gondii* forming a heteromeric complex (**Figure 2** and **Figure 3**), a hitherto undescribed phenomenon in nature.

There have been several unsuccessful attempts by us [unpublished] and others ^35–37^ to express a functional *Pf*PanK1. Whilst the protein has been successfully expressed in soluble form using various heterologous expression systems (*E. coli*, insect cells, *S. cerevisiae*), no study has reported PanK activity from the heterologously-expressed and purified protein. Our observation here that the presence of *Pf*PanK2 (as well as, potentially, additional proteins) is inextricably linked to PanK activity probably explains why these previous attempts of expressing an active *Pf*PanK1 were unsuccessful ^35–37^.

A comparison of the amino acid sequences of *P. falciparum* and *T. gondii* PanKs with those of other type II PanKs, such as human PanK3 (*Hs*PanK3), provides a possible explanation for why PanKs from these apicomplexan parasites exist in heteromeric complexes **(Figure S6)**. Each of the two identical active sites of the homodimeric *Hs*PanK3 are formed by parts of both of its protomers. Certain residues form hydrogen bonds with pantothenate (Glu138, Ser195, Arg207 from one protomer and Val268’ and Ala269’ from the second protomer), while others interact to stabilise the active site (Asp137 with Tyr258’, and Glu138 with Tyr254’) ^38, 39^ **(Figure S10)**. Notably, the hydrogen bond between Glu138 and Tyr254’ is important for the allosteric activation of the enzyme ^39^. Critically, one of the important residues involved in active site stabilisation, Asp137, is only conserved in the PanK1 of *P. falciparum* and *T. gondii* but not their PanK2, while others, such as Tyr254’ and Tyr258’ are conserved in their PanK2 but not PanK1 (**Figure S6** and **S10**). This raises the possibility that PanK1 and PanK2 homodimers are likely not functional, and that only a heteromeric PanK1/PanK2 complex, with a single complete active site, can serve as a functional PanK enzyme in these apicomplexan parasites. This is consistent with the previous observation that two of the nucleotide-binding motifs of *Pf*PanK2 deviate from those of other eukaryotic PanKs ^28^. Whether the incomplete second active site plays an additional, as yet undetermined, role(s) remains to be seen. It should be noted that the PanKs of other apicomplexan parasites exhibit a similar conservation of residues as that described above for *P. falciparum* and *T. gondii* (**Figure S11**), raising the possibility that heteromeric PanK complexes are ubiquitous in Apicomplexa.

The apparent molecular weight of the *Pf*PanK heterodimer complex (as determined from native western blotting) is consistent with that of a complex that includes *Pf*PanK1, *Pf*PanK2 and a *Pf*14-3-3I dimer (**Figure 1b**). However, due to various limitations of native gels ^40^, it is difficult to obtain an accurate estimate of the molecular weight of the complex. Although we cannot completely rule out the inclusion of other proteins in the *Pf*PanK complex, such as M17 leucyl aminopeptidase (**Figure 2a** and **Table S3**), we think that this is unlikely, since peptides from these proteins were detected at lower abundance than peptide from *Pf*PanK1, *Pf*PanK2 and *Pf*14-3-3I. The role of *Pf*14-3-3I in the heteromeric *Pf*PanK complex (**Figure 2a** and **2c**) is not clear. The 14-3-3 protein family comprises highly conserved proteins that occur in a wide array of eukaryotic organisms, including apicomplexans such as *P. falciparum* ^32, 41–43^. Multiple isoforms of 14-3-3 are found to occur in every organism that expresses the protein ^44^. 14-3-3 proteins bind to, and regulate, the function of proteins that are involved in a large range of cellular functions, including cell cycle regulation, signal transduction and apoptosis (reviewed in ^45^). They typically bind to phosphorylated Ser/Thr residues on target proteins, and modify their target protein’s trafficking/targeting (reviewed in ^46^), conformation, co-localisation, and/or activity (reviewed in ^47^). We speculate that *Pf*14-3-3I plays a regulatory role in the *Pf*PanK complex. The *Tg*PanK heterodimer complex has a molecular weight that is much larger than the combined molecular weights of *Tg*PanK1 and *Tg*PanK2. Unfortunately, mass spectrometry analysis of the *T. gondii* PanK complex was unsuccessful, presumably because the native level of expression of the complex is too low.

In this study, we have characterised, for the first time, PanK activity in *T. gondii*. The [^14^C]pantothenate phosphorylation data generated with the purified *Tg*PanK complex (**Figure 3c**) provide the first biochemical evidence indicating that these putative PanKs are able to phosphorylate pantothenate. This finding, combined with the results of the knockdown and *Sa*PanK complementation experiment in *T. gondii* (**Figure 4b** and **4c**), not only demonstrate the essentiality of *Tg*PanK1 and *Tg*PanK2, but also show that the essentiality is due to their role in phosphorylating pantothenate.

*T. gondii* parasites inhabit metabolically active mammalian cells that contain their own CoA biosynthesis pathway. Our data indicate that *T. gondii* parasites are unable to scavenge sufficient downstream intermediates in the CoA biosynthesis pathway from their host cells, including CoA, for their survival, and therefore must maintain their own active CoA biosynthesis pathway. The requirement for CoA biosynthesis in *T. gondii*, coupled with the intense investigation of this pathway as a drug target in *P. falciparum* ^27, 37, 48–61^, suggests that further characterisation of *Tg*PanK, and the CoA biosynthesis pathway in *T. gondii*, could yield novel drug targets for chemotherapy.

It has been an open question as to why many organisms (eukaryotes ^11–21^ and prokaryotes ^7, 23–25^), including all apicomplexan parasites ^62^, express more than one PanK and why some PanKs appear to be non-functional ^7, 21, 23^ (either by analysis of their sequence or through failed attempts to demonstrate PanK activity experimentally). The data that we present here provides a possible answer to this question.

## METHODS

### Parasite and host cell culture

*P. falciparum* parasites were maintained in RPMI 1640 media supplemented with 11 mM glucose (to a final concentration of 22 mM), 200 µM hypoxanthine, 24 µg/mL gentamicin and 6 g/L Albumax II as described previously ^63^. *T. gondii* was cultured in human foreskin fibroblasts (HFF cells) as described previously ^64^. *T. gondii* parasites were grown in flasks with a confluent HFF cell layer in either Dulbecco’s modified Eagle’s medium (DMEM) or complete RPMI 1640, with both media containing 2 g/L sodium bicarbonate and supplemented with 1% (v/v) fetal bovine serum (FBS), 50 units/mL penicillin, 50 µg/mL streptomycin, 10 µg/mL gentamicin, 0.2 mM L-glutamine, and 0.25 µg/mL amphotericin B.

### Plasmid preparation and parasite transfection

The *Pf*PanK1-GFP-expressing cell line was generated in a previous study ^27^, while the <u>untagged</u> GFP line was a generous gift from Professor Alex Maier (Research School of Biology, Australian National University, Canberra). A *Pfpank2*-pGlux-1 vector was generated for the overexpression of *Pf*PanK2-GFP in 3D7 strain *P. falciparum* as detailed in the **SI**. The primers used are listed in **Table S1**. The same construct was also transfected into each of the mutant clones and their Parent line described previously by Tjhin *et al*. ^27^. Transfections were performed with ring-stage parasites and transformants were subsequently selected and maintained using WR99210 (10 nM) as described previously ^65^.

The *Tg*PanK1-mAIDHA and *Tg*PanK2-mAIDHA expressing lines were generated using a CRISPR/Cas9 strategy as previously described in Shen *et al.* ^66^, which is detailed in the **SI**. The guide RNAs, primers, and the sequences of gBlocks used are provided in **Table S1** **and S2.**

The complementation lines *Tg*PanK1-mAIDHA^+*Sa*PanK-Ty1^ and *Tg*PanK2-mAIDHA^+*Sa*PanK-Ty1^ were created by expressing the *S. aureus* type II PanK gene (*Sapank*) in *T. gondii* under the regulation of the tubulin promoter (details in the **SI** and **Table S1** **and S2).**

### Immunofluorescence assays and microscopy

Fixed *Pf*PanK2-GFP-expressing 3D7 strain *P. falciparum* parasites within infected erythrocytes were observed and imaged with a Leica TCS-SP2-UV confocal microscope (Leica Microsystems) using a 63 × water immersion lens as described in the **SI**. To confirm the expression of *Sa*PanK-Ty1 in the *Tg*PanK1-mAIDHA^+*Sa*PanK-Ty1^ line, immunofluorescence assays were performed based on the protocol described by van Dooren *et al.* ^67^. *T. gondii* parasites were incubated with mouse anti-Ty1 antibodies (1:200 dilution). Secondary antibodies used were goat anti-mouse AlexaFluor 488 at a 1:250 dilution. The nucleus was stained with DAPI. Immunofluorescence images were acquired on a DeltaVision Elite system (GE Healthcare) using an inverted Olympus IX71 microscope with a 100 × UPlanSApo oil immersion lens (Olympus) paired with a Photometrics CoolSNAP HQ^2^ camera. Images taken on the DeltaVision setup were deconvolved using SoftWoRx Suite 2.0 software. Images were adjusted linearly for contrast and brightness.

### Polyacrylamide gel electrophoresis and western blotting

Parasite samples were analysed using either denaturing or blue native gels to determine the presence and abundance of a single protein or protein complex of interest, respectively. Briefly, mature trophozoite-stage *P. falciparum* parasites were isolated from infected erythrocytes by saponin lysis, as described previously ^68^. Saponin-isolated parasites were then pelleted and lysed in the appropriate buffers (as detailed in the **SI**). *T. gondii* protein samples were prepared as described previously, with samples for blue native-PAGE solubilised in Native PAGE sample buffer (ThermoFisher) containing 1% (v/v) Triton X-100^67^. Protein samples generated from both *P. falciparum* and *T. gondii* parasites were separated by polyacrylamide gel electrophoresis (PAGE) in precast NuPAGE (4-12% or 12%) or NativePAGE (4-16%) gels (ThermoFisher) according to the manufacturer’s instructions with minor modifications (detailed in the **SI**). The separated proteins were transferred to the appropriate membranes (nitrocellulose or polyvinylidene fluoride (PVDF)) and blocked (detailed in the **SI**) before immunoblotting. Blocked membranes were exposed (45 min – 2 h) to specific primary and secondary antibodies to allow for the detection of the protein of interest. To visualise the protein band(s), membranes were incubated in Pierce enhanced chemiluminescence (ECL) Plus Substrate (ThermoFisher) according to the manufacturer’s instructions or home-made ECL (0.04% w/v luminol, 0.007% w/v coumaric acid, 0.01% v/v H2O2, 100 mM Tris, pH 9.35). Protein bands were then either imaged onto X-ray films and scanned or visualised on a ChemiDoc MP Imaging System (Thermo Scientific).

### Flow cytometry

Saponin-isolated mature trophozoites from 3D7 wild-type, Parent^+*Pf*PanK2-GFP^, PanOH-A^+*Pf*PanK2-GFP^, PanOH-B^+*Pf*PanK2-GFP^ and CJ-A^+*Pf*PanK2-GFP^ cultures were subjected to flow cytometry analysis to determine the proportion of GFP-positive cells. Aliquots of each isolated parasite suspension were diluted in a saline solution (125 mM NaCl, 5 mM KCl, 25 mM HEPES, 20 mM glucose and 1 mM MgCl_2_, pH 7.1) to a concentration of ∼10^6^ – 10^7^ cells/mL in 1.2 mL Costar polypropylene cluster tubes (Corning) and sampled for flow cytometry analysis (in measurements of 100,000 cells, low sampling speed) with the following settings: forward scatter = 450 V (log scale), side scatter = 350 V (log scale) and AlexaFluor 488 = 600 V (log scale). The 3D7 wild-type cells were used to establish a gating strategy that defined a threshold below which parasites were deemed to be auto-fluorescent. This strategy was then applied in all analyses to determine the proportion of cells in each cell line that was GFP-positive (i.e. above the defined threshold).

### Immunoprecipitations

In order to immunopurify GFP-tagged or HA-tagged proteins from parasite lysates, immunoprecipitation was performed using either GFP-Trap (high affinity anti-GFP alpaca nanobody bound to agarose beads; Chromotek) or anti-HA beads (Sigma-Aldrich), respectively. *P. falciparum* lysate was prepared from saponin-isolated trophozoites, and *T. gondii* lysate was prepared from tachyzoites, as described previously (^68^ and ^67^, respectively). Immunoprecipitation was then performed (as detailed in the **SI**). In *P. falciparum* experiments where the amount of immunoprecipitated proteins were to be standardised across cell lines and biological repeats, the number of GFP-positive cells to be used for lysate preparation was calculated by a combination of haemocytometer count and flow cytometry. All immunoprecipitated samples from Parent^+*Pf*PanK2-GFP^, PanOH-A^+*Pf*PanK2-GFP^, PanOH-B^+*Pf*PanK2-GFP^ and CJ-A^+*Pf*PanK2-GFP^ cell lines contained protein from 5 × 10^7^ GFP-positive cells. Each of these samples were subsequently divided into two equal aliquots, one used in the [^14^C]pantothenate phosphorylation assay and the other for denaturing western blot.

When an aliquot of the immunoprecipitation sample (beads that have bound proteins from ∼10^6^ – 10^7^ GFP-positive cells for *P. falciparum* and ∼10^7^ – 10^8^ cells for *T. gondii*) was required for western blot, the bead suspension was centrifuged (2,500 × *g*, 2 min), the supernatant removed, and the beads resuspended in 50 µL sample buffer containing 2 × NuPAGE lithium dodecyl sulfate (LDS) sample buffer (ThermoFisher) and 2 × NuPAGE sample reducing agent (ThermoFisher). In some experiments, 10 µL aliquots of the total and unbound lysate fractions were each mixed with 10 µL of the same sample buffer. These samples were then boiled (95 °C, 10 min) and 10 µL of each was then used in a denaturing western blot as described above.

### [^14^C]Pantothenate phosphorylation assays

In order to determine the PanK activity of the protein(s) isolated in the GFP-Trap immunoprecipitation assays, the immunopurified complexes were used to perform a [^14^C]pantothenate phosphorylation time course. The bead suspensions containing the immunoprecipitated proteins from *P. falciparum* and *T. gondii* were centrifuged (2,500 × *g*, 2 min), the supernatant removed, and the beads resuspended in 300 µL (for *T. gondii*) or 500 µL (for *P. falciparum*) of buffer containing 100 mM tris(hydroxymethyl)aminomethane (Tris)-HCl (pH 7.4), 10 mM ATP and 10 mM MgCl_2_ (i.e. all reagents were at twice the final concentration required for the phosphorylation reaction). Each time course was then initiated by the addition of 300 µL (for *T. gondii*) or 500 µL (for *P. falciparum*) of 4 µM (0.2 µCi/mL) [^14^C]pantothenate in water (pre-warmed to 37 °C), to the bead suspension. Aliquots of each reaction (50 µL in duplicate) were terminated at pre-determined time points by mixing with 50 µL 150 mM barium hydroxide preloaded within the wells of a 96-well, 0.2 µm hydrophilic PVDF membrane filter bottom plate (Corning). Phosphorylated compounds in each well were then precipitated by the addition of 50 µL 150 mM zinc sulfate to generate the Somogyi reagent ^69^, the wells processed, and the radioactivity in the plate determined as detailed previously ^70^. Total radioactivity in each phosphorylation reaction was determined by mixing 50 µL aliquots of each reaction (in duplicate) thoroughly with 150 µL Microscint-40 (PerkinElmer) by pipetting the mixture at least 50 times, in the wells of an OptiPlate-96 microplate (PerkinElmer) ^70^.

### Mass spectrometry of immunoprecipitated samples

The identities of the proteins co-immunoprecipitated from lysates of parasites expressing *Pf*PanK1-GFP, *Pf*PanK2-GFP and untagged GFP were determined by mass spectrometry. Aliquots of bead-bound co-immunoprecipitated samples were resuspended in 2 × NuPAGE LDS sample buffer and 2 × NuPAGE sample reducing agent and sent (at ambient temperature, travel time less than 24 h) to the Australian Proteomics Analysis Facility (Sydney) for processing and mass spectrometry analysis (as detailed in the **SI**).

### Fluorescent *T. gondii* proliferation assay

Fluorescent *T. gondii* proliferation assays were performed as previously described ^34^. Briefly, 2000 parasites suspended in complete RPMI were added to the wells of optical bottom black 96 well plates (ThermoFisher) containing a confluent layer of HFF cells, in the presence or absence of 100 µM IAA, in triplicate. Fluorescent measurements (Excitation filter, 540 nm; Emission filter, 590 nm) using a FLUOstar OPTIMA Microplate Reader (BMG LABTECH) were taken over 7 days.

### Knockdown of mAID protein

Flasks containing a confluent layer of HFF cells were seeded with *Tg*PanK1-mAIDHA, *Tg*PanK2-mAIDHA, *Tg*PanK1-mAIDHA^+*Sa*PanK-Ty1^ or *Tg*PanK2-mAIDHA^+*Sa*PanK-Ty1^ *T. gondii* parasites. While the parasites were still intracellular, 100 µM of IAA dissolved in ethanol (final ethanol concentration of 0.1%, v/v) was added to induce the knockdown of *Tg*PanK1-mAIDHA or *Tg*PanK2-mAIDHA, with ethanol (0.1%, v/v) added to another flask as a vehicle control. Flasks with IAA added were processed at 1, 2 and 4 h time points, and the control flask was processed at the 4-hour time point. Parasite concentrations were determined using a haemocytometer and 1.5 × 10^7^ parasites were resuspended in LDS sample buffer, and boiled at 95 °C for 10 minutes. An aliquot from each sample was analysed by western blotting.

### Alignment of PanK

PanK homologues from *P. falciparum* and *T. gondii*, and a selection of other type II PanKs from other eukaryotic organisms and *S. aureus* were aligned using PROMALS3D ^71^ (available at: http://prodata.swmed.edu/promals3d/promals3d.php). The default parameters were selected except for the ‘Identity threshold above which fast alignment is applied’ parameter, which was changed to “1” to allow for a more accurate alignment.

## ACKNOWLEDGEMENTS

Proteomics was undertaken at APAF, the infrastructure provided by the Australian Government through the National Collaborative Research Infrastructure Strategy (NCRIS). We are grateful to the Canberra Branch of the Australian Red Cross Blood Service for the provision of red blood cells, and Professor Alex Maier (ANU) for the untagged GFP-expressing *P. falciparum* parasites. ETT, VMH and CS were supported by Research Training Program scholarships from the Australian Government. CS was also funded by an NHMRC Overseas Biomedical Fellowship (1016357). This work was, in part, supported by a Project Grant (APP1129843) from the National Health and Medical Research Council to KJS and a Discovery Grant (DP150102883) from the Australian Research Council to GvD.

The authors declare that they have no conflict of interest.

## Supplementary Information

### METHODS

#### Plasmid preparation

The *Pfpank2*-pGlux-1 construct has the *Pfpank2*-coding sequence inserted within multiple cloning site (MCS) III of pGlux-1. The plasmid backbone contains the human dihydrofolate reductase (*hdhfr*) gene, which confers resistance to WR99210, as a positive selectable marker. *Pfpank2* is placed under the regulation of the *Plasmodium falciparum* chloroquine resistance transporter (*Pfcrt*) promoter, and upstream of the GFP-coding sequence.

The *Pfpank2* sequence used to generate the *Pfpank2*-pGlux-1 construct was initially amplified from parasite cDNA. Total RNA was purified from saponin-isolated *P. falciparum* parasites (typically 2 × 10^7^ cells) using the RNeasy Mini Kit (QIAGEN) according to the manufacturer’s protocol for purifying total RNA from animal cells. The optional 15 min DNase I incubation was included to eliminate residual genomic DNA. Complementary DNA (cDNA) was synthesised from this total RNA sample using SuperScript II Reverse Transcriptase (ThermoFisher) with Oligo(dT)_12-18_ primer and included the optional incubation with RNaseOUT Recombinant Ribonuclease Inhibitor (ThermoFisher), all according to the manufacturer’s protocol. The *Pfpank2*-specific sequence was then amplified from cDNA using Platinum *Pfx* DNA polymerase (ThermoFisher) with the oligonucleotide primers listed in **Table S1**. The *Pfpank2*-coding sequence was then inserted into pGlux-1 using the In-Fusion cloning (Clontech) method. Before cloning, the pGlux-1 plasmid was linearised by sequential digestions with *Xho*I (ThermoFisher) and subsequently *Kpn*I (New England Biolabs), according to each manufacturer’s recommendation. The In-Fusion reaction was set up with the In-Fusion Dry-Down PCR Cloning Kit, essentially as described in the manufacturer’s protocol.

The *Tg*PanK1-mAIDHA and *Tg*Pank2-mAIDHA expressing lines were generated using a CRISPR/Cas9 based genome editing approach as previously described in Shen *et al.*^1^. Guide RNA (gRNA) sequences that would enable Cas9 to complex with the RNA to cut close to the 3’ end of either the *Tgpank1* or *Tgpank2* gene were incorporated into pSAG1::Cas9-U6::sgUPRT (Addgene plasmid #54467 ^1^) by utilising the Q5 site-directed mutagenesis kit (New England Biolabs). This included performing an initial PCR incorporating the gRNA into the vector and subsequently circularising the vector. This vector encodes both the gRNA and the Cas9-GFP. A gBlock (IDT) containing the mAIDHA construct ^2^ was also amplified using gene-specific primers with homologous ends to the locus targeted by the gRNA to enable homologous recombination. The gRNA-expressing pSAG1::CAS9-U6::sgUPRT construct for each *Tgpank* gene was transfected with the corresponding mAIDHA gBlock fragment into RH strain TATiΔKu80:TIR1 tachyzoite-stage parasites ^2^ also expressing a tandem dimeric (td) Tomato red fluorescent protein. The GFP expression from the Cas9-GFP enabled the use of flow cytometry to select GFP positive cells to establish a clonal population two days after transfection. Transformants were identified by PCR screening (all primers and gBlocks are listed in **Table S1**).

The *Tg*PanK1-GFP/*Tg*PanK2-mAIDHA expressing line was generated utilising the CRISPR/Cas9 approach as mentioned above. A gBlock (IDT) containing the TEV-GFP construct was amplified using gene-specific primers with homologous ends to the *Tgpank1* locus targeted by the gRNA to enable homologous recombination. This was transfected with the gRNA-expressing pSAG1::CAS9-U6::sgUPRT construct for *Tgpank1* into the *Tg*PanK2-mAIDHA line that had been previously created. GFP positive cells were selected for as noted above and a clonal line was confirmed by PCR screening (all primers and gBlock are listed in **Table S1**). The *Tg*PanK1-HA/*Tg*PanK2-GFP expressing line was generated in RH strain TATiΔKu80 parasites using the same CRISPR/Cas9 approach with the TEV-HA gBlock and TEV-GFP gBlock (IDT).

The complementation lines *Tg*PanK1-mAIDHA^+*Sa*PanK-Ty1^ and *Tg*PanK2-mAIDHA^+*Sa*PanK-Ty1^ were created by expressing Ty1-tagged *Staphylococcus aureus* type II pantothenate kinase (*Sapank*) in the *Tg*PanK1-mAIDHA and *Tg*PanK2-mAIDHA *T. gondii* strains. Briefly, the *Sapank* open reading frame was PCR amplified using a gBlock encoding *Sapank* that had been codon-optimised for *T. gondii* expression. The resultant PCR product was digested with *Bgl*II and *Avr*II and ligated into the equivalent sites of the vector pUBTTY. The pUBTTy was modified from the vector pBTTy, described previously ^3^, which contains a phleomycin resistance marker, and an expression cassette containing the *T. gondii α*-tubulin promoter and a Ty1 epitope tag. The UPRT flank was digested from the vector pUgCTH3 ^4^ with *Apa*I and *Hin*dIII and ligated into the equivalent sites of pBTTy to generate pUBTTy. The *Sa*pank-Ty1 containing pUBTTy vector was linearised and transfected into the lines expressing *Tg*PanK1-mAIDHA and *Tg*PanK2-mAIDHA. Transformants were subsequently selected using phleomycin (50 µg/mL) in DMEM supplemented with 10 mM Hepes and 10 µg/mL gentamicin, pH 7.6, for 4 h as described previously ^5^. Phelomycin-resistant parasites were cloned using fluorescence activated cell sorting, and subsequently cultured in complete RPMI-1640.

#### Preparation of cells for microscopy

Coverslip-bound *P. falciparum* infected red blood cells were first prepared by washing (500 × *g*, 5 min) parasite-infected erythrocytes (5 − 10% parasitaemia) once and resuspending them at ∼2% haematocrit in 137 mM NaCl, 2.7 mM KCl, 10 mM phosphate buffer, pH 7.4 (phosphate buffered saline; PBS). Next, 1 − 2 mL of the suspension was added to a polyethylenimine (PEI)-coated coverslip placed within a well of a 6-well plate. Plates were incubated (with shaking) for 15 min at room temperature and unbound cells were subsequently washed off the coverslips with PBS (2 mL per well, with a 2 min shaking incubation followed by aspiration). Cells were then fixed with 1 mL of PBS containing 4% (w/v) paraformaldehyde (Electron Microscopy Services) and 0.0075% (w/v) glutaraldehyde (30 min at room temperature). The fixative was then aspirated, and the coverslips washed in PBS three times as described above, before they were rinsed in water and dried. A drop of SLOWFADE (Invitrogen) containing the nuclear stain 4’,6-diamidino-2-phenylindole (DAPI) was added to the centre of the coverslips. Finally, each coverslip was inverted onto a microscope slide, sealed with nail polish and used for confocal imaging.

For *T. gondii* immunofluorescence assays, parasites were inoculated onto HFF-coated coverslips and allowed to proliferate overnight. Parasites were fixed in 3% (w/v) paraformaldehyde in PBS, permeabilised in 0.25% (v/v) Triton X-100 in PBS and blocked in 2% (w/v) bovine serum albumin in PBS. Parasites were incubated in mouse anti-Ty1 primary antibodies (1:200 dilution; ^6^) and goat anti-mouse AlexaFluor 488-conjugated secondary antibodies (1:250 dilution; ThermoFisher, catalogue number A11029).

#### Denaturing polyacrylamide gel electrophoresis

Saponin-isolated *P. falciparum* parasites (typically ∼10^8^ cells) were centrifuged (15,850 × *g*, 30 s) and the supernatant was removed. The pellet was resuspended in 200 µL of lysis buffer (1 × mini cOmplete protease inhibitor cocktail (Roche), 1 × NuPAGE LDS sample buffer (ThermoFisher), 1 × NuPAGE sample reducing agent (ThermoFisher), 50 – 60 units of benzonase nuclease (Novagen) and 7.5 – 10 mM of MgCl_2_), mixed well by vortexing and then incubated at 95 °C for 10 min. The sample was then centrifuged (16,000 × *g*, 30 min) to pellet the haemozoin before the supernatant was used for gel electrophoresis. Each sample (10 µL) was subsequently loaded into separate wells of a NuPAGE 4 – 12% Bis-Tris protein gel (1.0 mm, 12 wells; ThermoFisher) alongside 5 µL of SeeBlue Plus2 pre-stained protein standards (ThermoFisher).

*T. gondii* tachyzoites, either freshly egressed from host HFF cells or mechanically egressed through a 26-gauge needle, were filtered through a 3 µm polycarbonate filter. Tachyzoites (typically 1.5 × 10^7^ cells), were subsequently centrifuged (12,000 × *g*, 1 min). The supernatant was aspirated and the pellet was resuspended in 30 µL of 1 × NuPAGE LDS sample buffer (ThermoFisher). The sample was mixed well by vortexing and incubated at 95 °C for 10 min. Samples were then frozen at -20 °C or used immediately for gel electrophoresis. Each sample (10 – 20 µL) was subsequently loaded into separate wells of a NuPAGE 4 – 12% Bis-Tris protein gel (1.0 mm, 12 wells; ThermoFisher) alongside 5 µL of Novex Sharp Pre-Stained Protein Standard (Invitrogen). Where relevant, parasites were incubated in 100 µM idole-3-acetic acid (IAA) or in a 0.1% (v/v) ethanol vehicle control for specified times prior to sample preparation.

Electrophoresis of *P. falciparum* and *T. gondii* samples was performed in 1 × NuPAGE 2-(*N*-morpholino)ethanesulfonic acid (MES) sodium dodecyl sulfate (SDS) running buffer (ThermoFisher) at 200 V for 30 – 35 min. Separated proteins were transferred (35 V for 1.5 h or 30 V for 1 h depending on the transfer system) to a nitrocellulose membrane (ThermoFisher) in 1 × NuPAGE transfer buffer containing 10% (v/v) methanol. The membrane was then blocked in a solution containing 4% (w/v) skim milk powder in Tris buffered saline (TBS) or PBS with shaking, either overnight at 4 °C or for 1 – 2 h at room temperature.

The primary antibodies used in this study included mouse anti-GFP monoclonal antibody (0.4 µg/mL final concentration; Roche, Sigma catalogue 11814460001), rat anti-HA monoclonal antibody (1.6 µg/mL final concentration; Sigma, clone 3F10, catalogue A11867431001), mouse anti-Ty1 monoclonal antibody (1:1000 dilution; ^6^), and a pan-specific anti-14-3-3 rabbit polyclonal antibody (0.25 µg/mL final concentration; Abcam). The secondary antibodies used for the *P. falciparum* blots were a goat anti-mouse horseradish peroxidase (HRP)-conjugated antibody and a goat anti-rabbit HRP-conjugated antibody (both 0.08 µg/mL final concentration; Santa Cruz Biotechnology). The secondary antibodies used for *T. gondii* experiments were goat anti-mouse HRP-conjugated antibody (0.4 – 0.8 µg/mL final concentration; Abcam), goat anti-rabbit HRP-conjugated antibody (0.1 µg/mL final concentration; Abcam), and goat anti-rat HRP-conjugated antibody (0.1 µg/mL final concentration; Abcam).

All antibodies were diluted in 4% (w/v) skim milk in TBS or PBS. After each antibody incubation, membranes were washed at least three times (5 – 10 min each) in fresh 0.05 or 0.1% (v/v) Tween 20 in TBS or PBS.

#### Native polyacrylamide gel electrophoresis

In order to detect the presence and abundance of protein(s) of interest in their native conformation, parasite samples were subjected to blue native gel electrophoresis. Briefly, saponin-isolated trophozoite-stage *P. falciparum* parasites (4 – 8 × 10^8^ cells) were centrifuged (15,850 × *g*, 30 s) and the supernatant removed from the pellet. The parasites were then resuspended by vortexing in 200 µL of lysis buffer containing 1 × mini cOmplete protease inhibitor cocktail (Roche), 1 × NativePAGE sample buffer (ThermoFisher), 0.5% (w/v) digitonin, 2 mM EDTA, 50 – 60 units of benzonase nuclease (Novagen) and 7.5 – 10 mM of MgCl_2_, and incubated with tumbling end-over-end at 4 °C. The lysis preparation was then centrifuged at 16,000 × *g* for 30 min at 4 °C and the supernatant immediately used for gel electrophoresis. Prior to electrophoresis, NativePAGE 5% G-250 sample additive (ThermoFisher) was added to each sample supernatant to a final concentration of 0.125% (w/v) and mixed by vortexing. Samples (typically 10 µL) were then loaded into the wells of a NativePAGE 4 – 16% Bis-Tris protein gel (1.0 mm, 10 wells; ThermoFisher). NativeMark unstained protein standards (5 µL, ThermoFisher) were loaded into the gel alongside the samples to allow for protein mass determination. Electrophoresis was carried out at 4 °C according to the manufacturer’s protocol for detergent-containing samples to be used for western blotting. At the end of the run, the proteins within the gel were transferred as described for denaturing western blot above to a methanol-primed 0.45 µm PVDF membrane (GE healthcare). At the end of the transfer, proteins were fixed to the membrane by a 15 min incubation in 10% (v/v) acetic acid and briefly rinsed in water. In order to visualise the ladder, the membrane was very briefly (∼5 s) de-stained in absolute methanol and rinsed in water before it was blocked overnight in 4% (w/v) skim milk powder in PBS at 4 °C with shaking.

*T. gondii* tachyzoites freshly or mechanically egressed from their host HFF cells were filtered through a 3 µm polycarbonate filter. Tachyzoites (typically 1.5 × 10^7^ cells) were subsequently centrifuged (12,000 × *g*, 1 min). The supernatant was removed and the pellet was resuspended by vortexing in 30 µL of lysis buffer containing 1 × mini cOmplete protease inhibitor cocktail (Roche), 1 × NativePAGE sample buffer (ThermoFisher), 10% (v/v) Triton X-100, 2 mM EDTA, and incubated with intermittent perturbation on ice for 30 min. The lysate was then centrifuged at 20,000 × *g* for 30 min at 4 °C and the supernatant was either stored at -20 °C or used immediately for gel electrophoresis. Sample preparation, electrophoresis and transfer was carried out as described for *P. falciparum.* Where anti-HA blotting was required on a PVDF membrane, after electrophoresis of the samples the membrane was placed in TBS containing 0.05% (v/v) Tween 20 overnight, then blocked in 4% (w/v) skim milk powder in TBS at room temperature for a minimum of 1 h the next day before probing with the anti HA antibody.

#### Immunoprecipitation

Briefly, saponin-isolated *P. falciparum* trophozoites were resuspended in 500 µL of lysis buffer containing 1 × mini cOmplete protease inhibitor cocktail (Roche) or 1 × Halt protease inhibitor cocktail (EDTA-free; ThermoFisher), GFP-Trap wash buffer (10 mM Tris/Cl, pH 7.5, 150 mM NaCl and 0.5 mM EDTA), 0.5% (w/v) digitonin, 50 – 60 units of benzonase nuclease (Novagen) and 3 – 4 mM of MgCl_2_. The pellet was resuspended well by vortexing and the suspension was incubated (30 – 60 min) with tumbling end-over-end at 4 °C. Subsequently, the suspension was centrifuged (16,000 × *g*, 30 min, 4 °C) and the supernatant used for GFP-Trap binding. Prior to immunoprecipitation, 25 µL (for each lysate) of GFP-Trap-agarose bead slurry was primed by three washes (2,500 × *g*, 2 min, 4 °C) in 500 µL of GFP-Trap wash buffer. The supernatant was removed from the beads at the end of the third wash and 450 – 500 µL of the lysate generated in the parasite lysis step was applied to the beads. In some experiments, 50 µL of the total lysate was collected for western blotting. This suspension was then incubated for 1 h at 4 °C with tumbling end-over-end. At the end of the incubation, the bead suspension was centrifuged (2,500 × *g*, 2 min, 4 °C) and the supernatant was discarded. In some experiments, 50 µL of this supernatant was collected to be used in western blots as the unbound fraction. The proteins bound to the beads were washed 3 × (2,500 × *g*, 2 min, 4 °C) in GFP-Trap wash buffer with or without 1 × mini cOmplete protease inhibitor cocktail or 1 × Halt protease inhibitor cocktail (EDTA-free). After removing the supernatant at the end of the third wash, the beads were resuspended in GFP-Trap wash buffer (typically 200 – 300 µL) and aliquots of these were used for downstream experiments.

*T. gondii* GFP-tagged proteins were purified using the GFP-Trap approach mentioned above, and the HA-tagged proteins were purified using anti-HA Affinity Matrix (Roche) following the manufacturer’s instructions with some modifications. Egressed *T. gondii* tachyzoites were filtered through a 3 µm polycarbonate filter. Parasites (∼10^7^-10^8^ for each line) were centrifuged (1,500 × *g*, 10 min, 4 °C). The supernatant was aspirated, and the cells were resuspended in 1 mL of PBS and centrifuged (12,000 × *g*, 1 min, 4 °C). After aspiration, lysis buffer (1 mL) containing 1 × mini cOmplete protease inhibitor cocktail (Roche), wash buffer (10 mM Tris/Cl, pH 7.5, 150 mM NaCl and 0.5 mM EDTA), and 1% (v/v) Triton X-100 was added to the remaining cells. The pellet was resuspended and the suspension was incubated (1 h) with tumbling end-over-end at 4 °C. Subsequently, the suspension was centrifuged (21,000 × *g*, 30 min, 4 °C). Prior to immunoprecipitation, 25 µL of GFP-Trap-agarose bead slurry, or 25 µL of anti-HA Affinity Matrix were washed three times in 500 µL of wash buffer (with 1% (v/v) Triton X-100 in the anti-HA wash buffer), with centrifugation at 2,500 × *g*, 2 min, 4 °C between washes. The supernatant was removed from the beads after the third wash and 450 µL of the parasite lysate was applied to each of the beads. A 50 µL aliquot of the total lysate was collected for western blotting as the ‘total’ fraction. The lysate/bead suspension was incubated for at least 1 h at 4 °C with tumbling end-over-end. At the end of the incubation, the bead suspension was centrifuged (2,500 × *g*, 2 min, 4 °C), 50 µL of this supernatant was collected for use in western blots as the ‘unbound’ fraction. The proteins bound to the beads were washed 3 × in wash buffer with centrifugation at 2,500 × *g*, 2 min, 4 °C between washes. After removing the supernatant at the end of the third wash, the beads were resuspended in wash buffer (typically 100 – 300 µL) and aliquots of these were used for downstream experiments. Alternatively, 100 µL of 1 × NuPAGE LDS sample buffer (ThermoFisher) was added to the samples to elute proteins from the beads and generate the ‘bound’ fractions.

#### Mass spectrometry of immunoprecipitated samples

The immunoprecipitated proteins were processed and identified through mass spectrometry (MS) analysis at the Australian Proteomics Analysis Facility. First, the loading buffer in the samples were separated from the proteins through a short one-dimension gel electrophoresis. The samples were denatured at 95 °C for 10 min and 2 × 15 µL of each sample was loaded into the lanes of a 12% iGel protein gel (1.0 mm, 12-well; NuSep). The gel was run at 15 mA for 25 min and subsequently washed with a solution containing 10% (v/v) methanol and 7% (v/v) acetic acid for 15 min. The gel was then washed with a fixant for 90 min and stained overnight in Coomassie. The band corresponding to the proteins was subsequently excised and de-stained with ammonium bicarbonate/acetonitrile (ACN). The protein samples were then reduced with 25 mM dithiothreitol (DTT) at 60 °C for 30 min and alkylated with 55 mM iodoacetamide before an in-gel protein digestion was performed overnight using 200 ng trypsin. The peptides generated were extracted from the gel with bath sonication and ACN/formic acid (FA), dried and then reconstituted in 30 µL of loading buffer.

The peptide samples were then subjected to 1D nano liquid chromatography tandem mass spectrometry (Nano-LC-ESI MS/MS) analysis. Sample (10 µL) was injected onto a peptide trap (Halo C18, 150 µm × 5 cm) for pre-concentration and desalted with 0.1% (v/v) FA, 2% (v/v) ACN at 4 µL/min for 10 min. The peptide trap was then switched into line with the analytical column. Peptides were subsequently eluted from the column using a linear solvent gradient, with steps, from H2O:ACN (98:2; + 0.1%, v/v, FA) to H2O:CH3CN (2:98; + 0.1%, v/v, FA) with constant flow (600 nL/min) over an 80 min period. The liquid chromatography eluent was subjected to positive ion nanoflow electrospray MS analysis in an information-dependent acquisition mode (IDA). In the IDA mode, a time-of-flight MS survey scan was acquired (*m/z* 350-1500, 0.25 s), with twenty largest multiply charged ions (counts >150) in the survey scan sequentially subjected to MS/MS analysis. MS/MS spectra were accumulated for 100 ms (*m/z* 100 – 1500) with rolling collision energy.

The peptides from the MS analysis were identified by comparing their amino acid sequences against an annotated protein database for *P. falciparum* 3D7 strain (version 28; PlasmoDB) using the ProteinPilot Software (version 4.2; SCIEX) at a detection threshold of >1.30 (95.0% confidence).

#### Fluorescent parasite proliferation assay

Fluorescent parasite proliferation assays were performed as previously described ^4,7^. Black 96-well plates containing confluent HFF host cells were washed with PBS (× 2). Complete RPMI-1640 (100 µL) with or without 200 µM of IAA supplementation (2 × final concentration), or with 10 µM pyrimethamine (2 × final concentration) for the ‘no growth’ control, were added to the relevant wells. Fluorescent parasites (parental line, *Tg*PanK1-mAIDHA, *Tg*PanK2-mAIDHA, *Tg*PanK1-mAIDHA^+*Sa*PanK-Ty1^ or *Tg*PanK2-mAIDHA^+*Sa*PanK-Ty1^) were plated in each well (100 µL, 2000 parasites) in triplicate. Plates were incubated at 37 °C in a 5% CO2 humidified incubator. Fluorescent measurements (Excitation filter, 540nm; Emission filter, 590nm) were taken up to two times a day over 7 days with the FLUOstar OPTIMA Microplate Reader (BMG LABTECH), and the proliferation of the fluorescent parasites was measured over this time. Values for the no growth control were considered as background values and were subtracted from the experimental values during data processing.

#### Alignment of PanK

The following PanK type II homologues annotated by accession number were aligned: *Staphylococcus aureus* (Q2FWC7), *Saccharomyces cerevisiae* (Q04430), *Aspergillus nidulans* (O93921), *Homo sapiens* PanK1 (Q8TE04) PanK2 (Q9BZ23) PanK3 (Q9H999) PanK4 (Q9NVE7), *Arabidopsis thaliana* PanK1 (O80765) PanK2 (Q8L5Y9)*, Plasmodium falciparum* PanK1 (Q8ILP4) PanK2 (Q8IL92) and *Toxoplasma gondii* PanK1 (A0A125YTW9) PanK2 (V5B595). We utilised PROMALS3D ^8^ (available at: http://prodata.swmed.edu/promals3d/promals3d.php) for the alignment.

#### Function

**Table S1.**
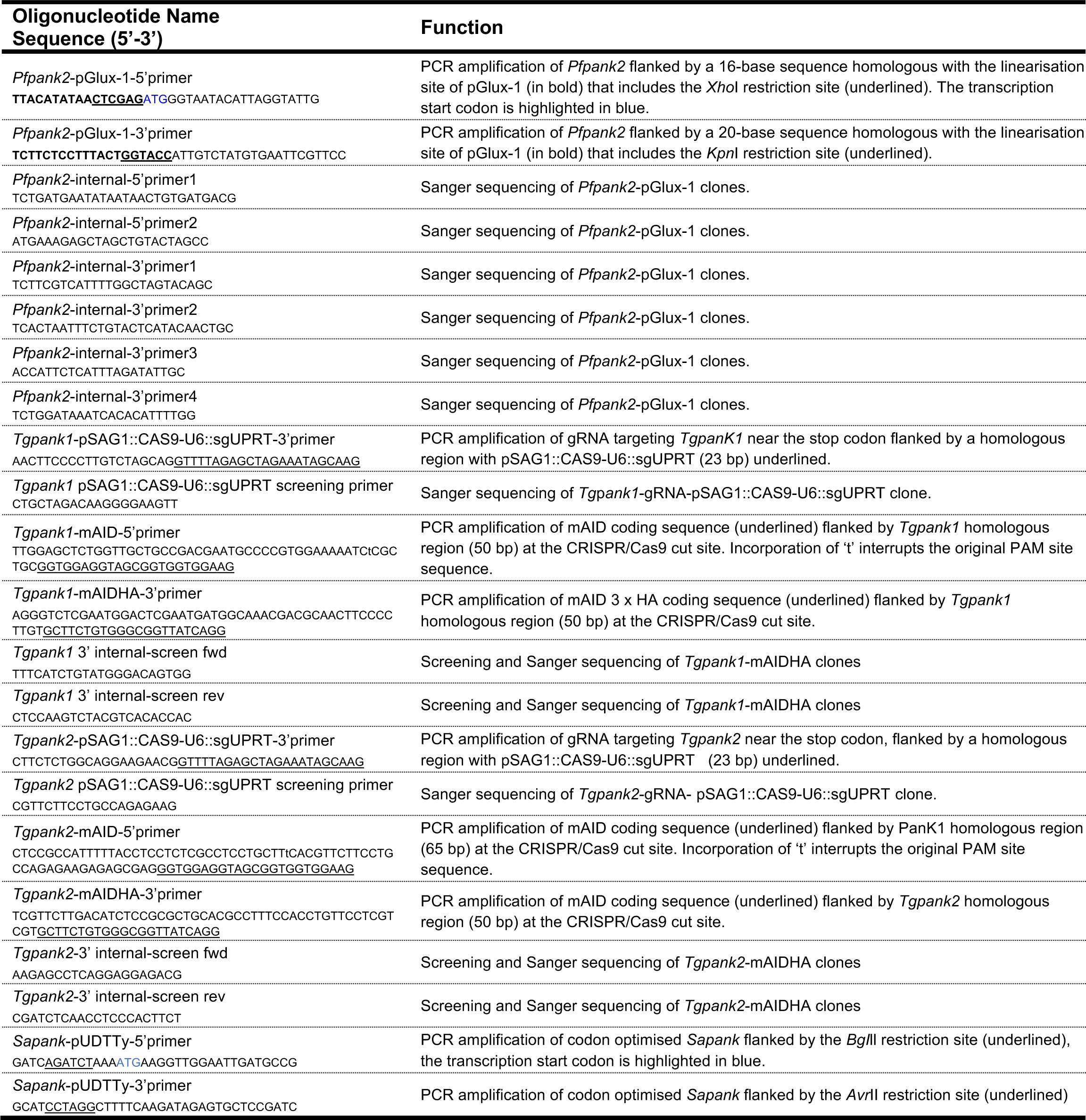
List of oligonucleotides used in this study.

**Table S2.**
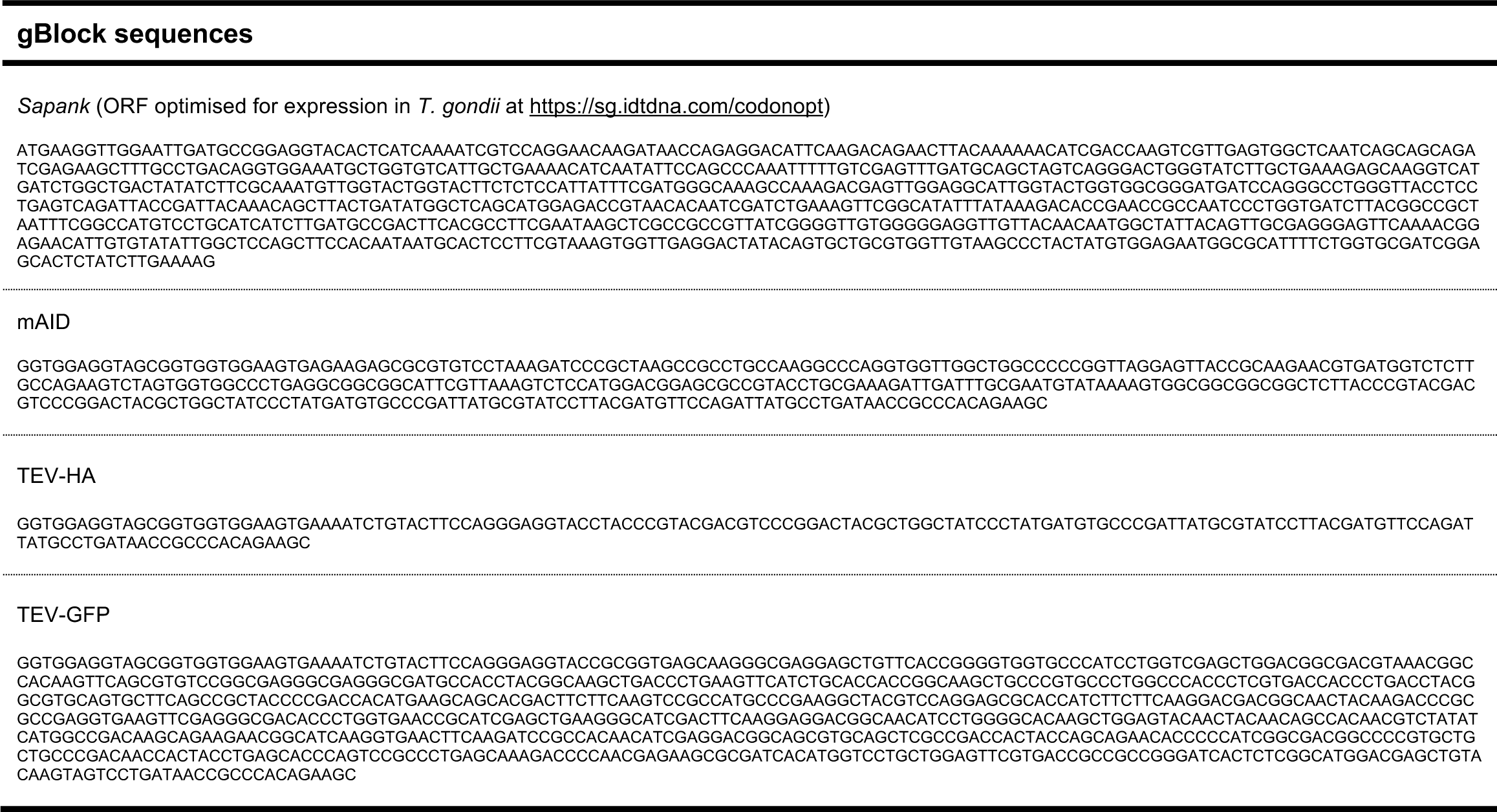
List of gBLOCK sequences used in this study.

**Table S3.**
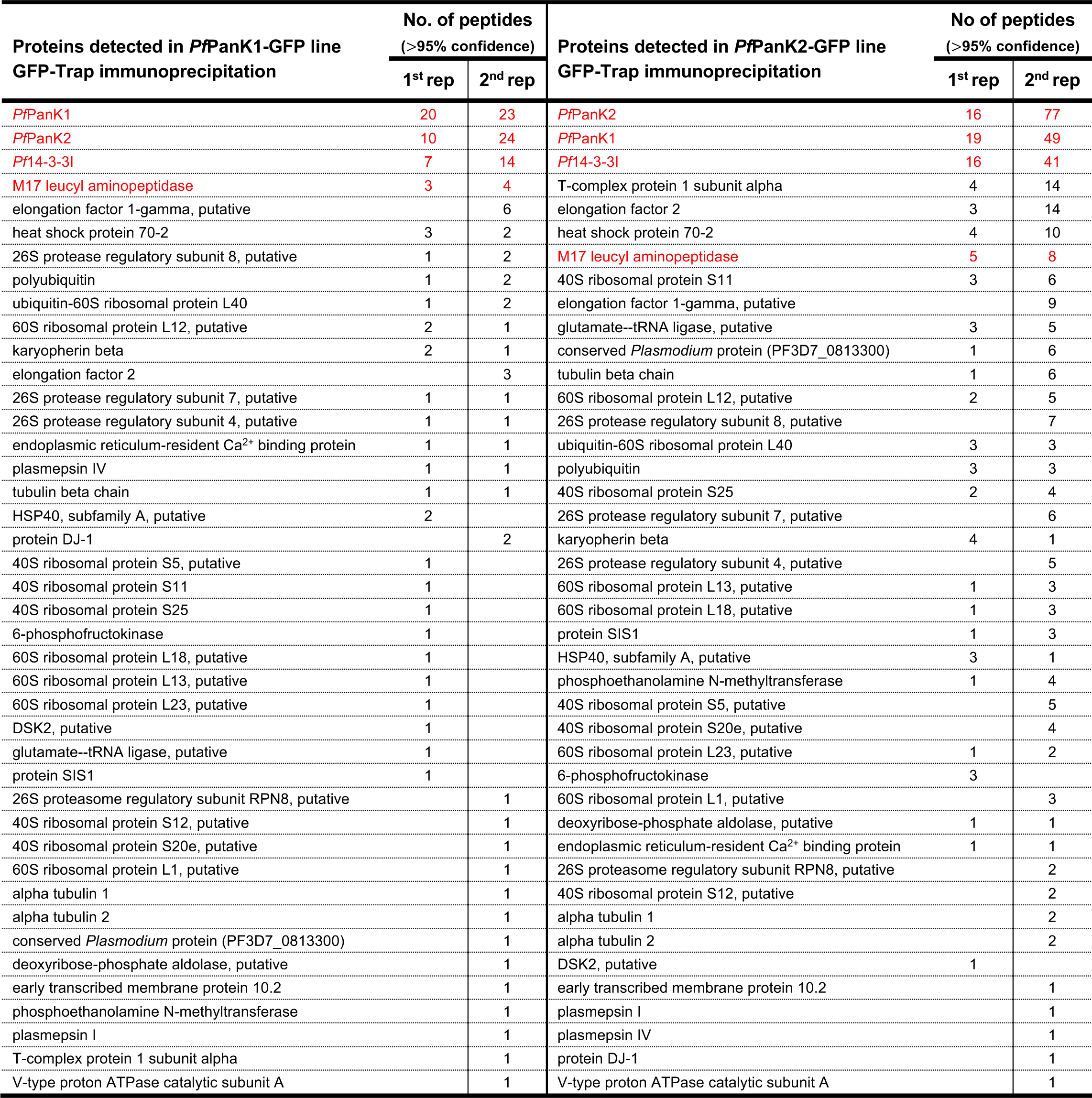
List of proteins identified in the MS analysis of the GFP-Trap immunoprecipitated complexes of *Pf*PanK1-GFP- and *Pf*PanK2-GFP-expressing parasites. Proteins are listed by the total number of peptides detected in the two independent replicates, from the most abundant to the least abundant. Only proteins that are present in the immunoprecipitation fractions of both parasite lines and absent in the negative controls (bound fractions of untagged GFP-expressing and 3D7 parasite lysates) are shown. Proteins shown in **Figure 2a** are indicated in red.

**Figure S1.**
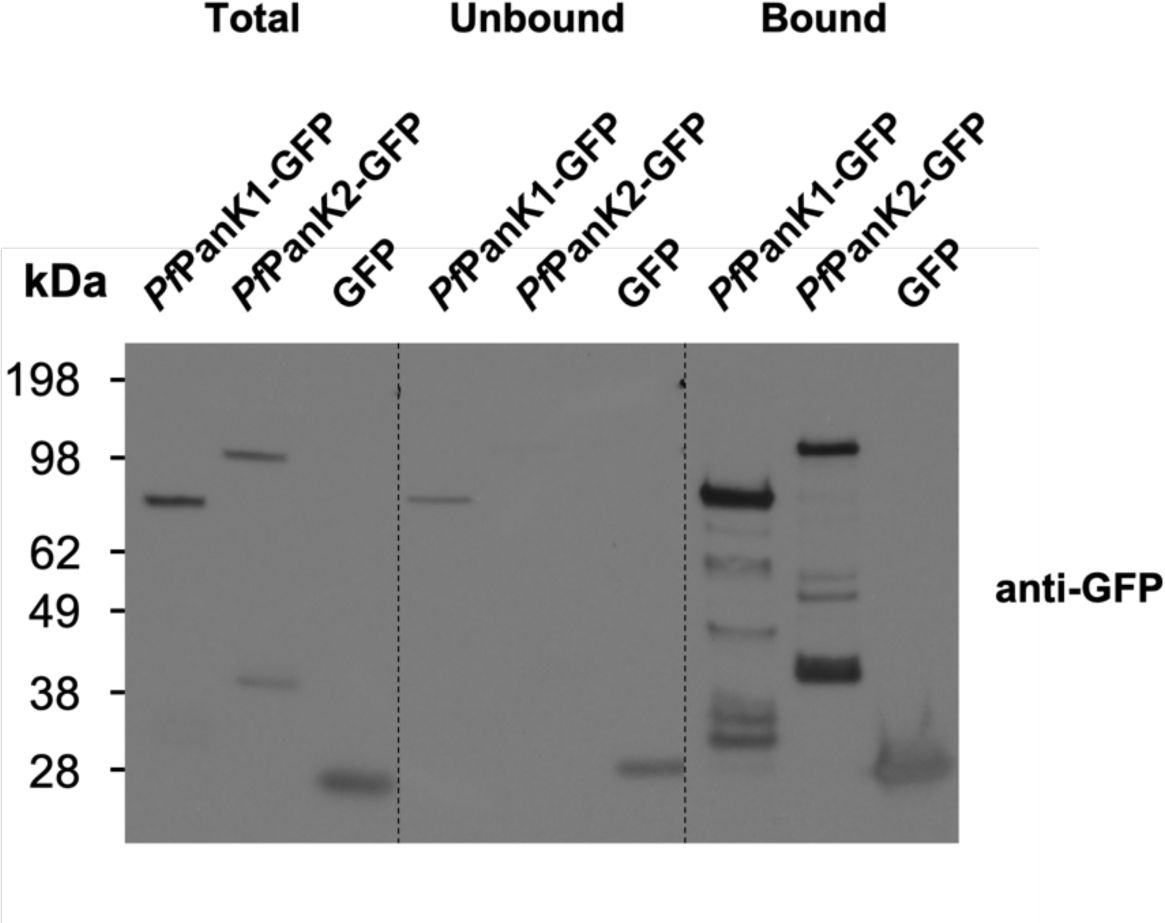
GFP-Trap immunoprecipitation of *Pf*PanK1-GFP-, *Pf*PanK2-GFP- and GFP-expressing parasites. Denaturing western blot analysis of the GFP-tagged proteins present in the total lysate, unbound and GFP-Trap-bound fractions of *Pf*PanK1-GFP-, *Pf*PanK2-GFP- and untagged GFP-expressing parasites. Western blots were performed with anti-GFP antibodies and the blot shown is representative of two independent experiments each performed with a different batch of parasites.

**Figure S2.**
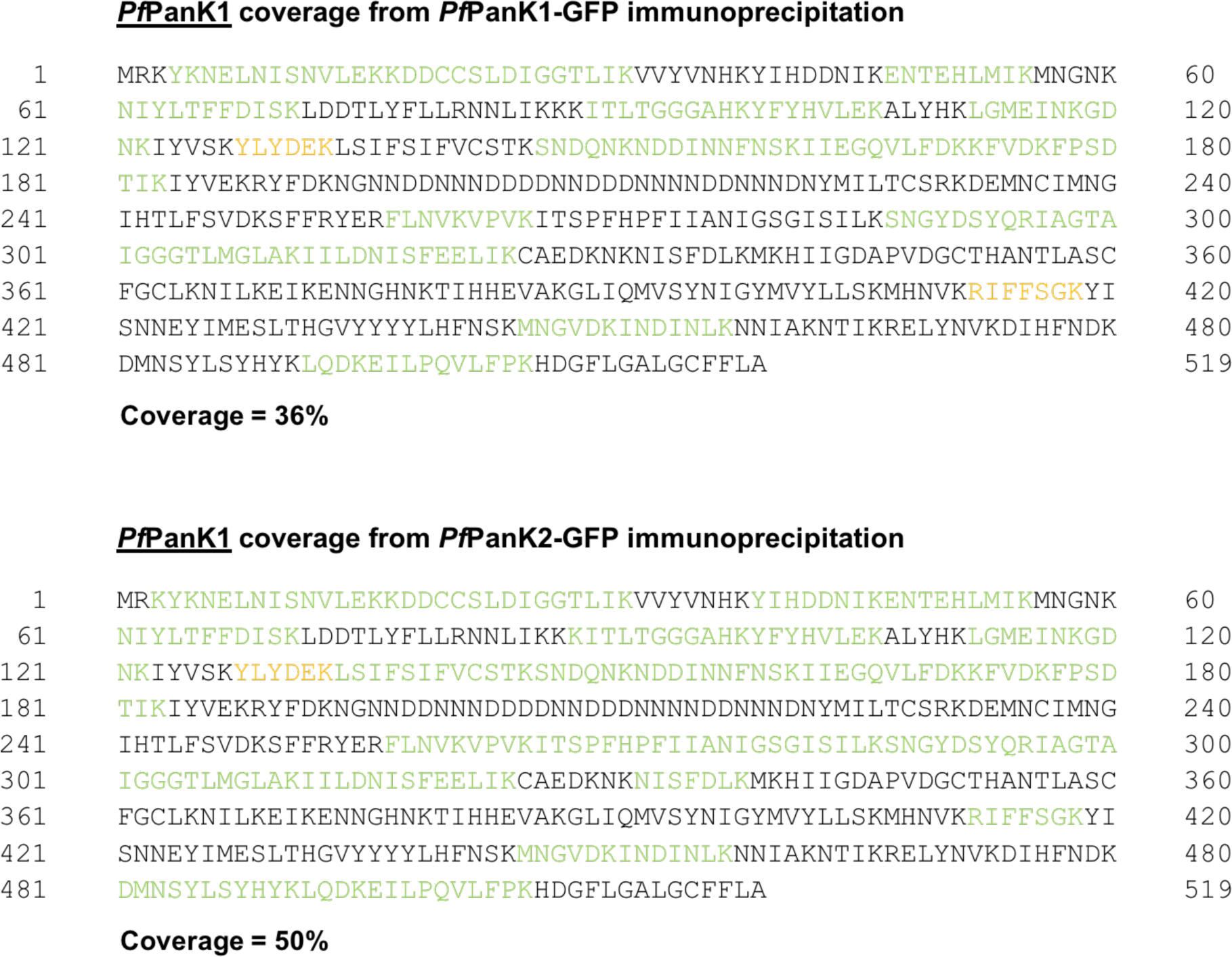
MS coverage of *Pf*PanK1. *Pf*PanK1 peptides detected in the two independent MS analyses of the GFP-Trap immunoprecipitation of the *Pf*PanK1-GFP- and *Pf*PanK2-GFP-expressing parasites. Residues in green were detected in either analysis with >95% confidence, while residues in orange were detected in either analysis with >90% (but <95%) confidence. Percentage coverage was calculated using only the residues labelled green.

**Figure S3.**
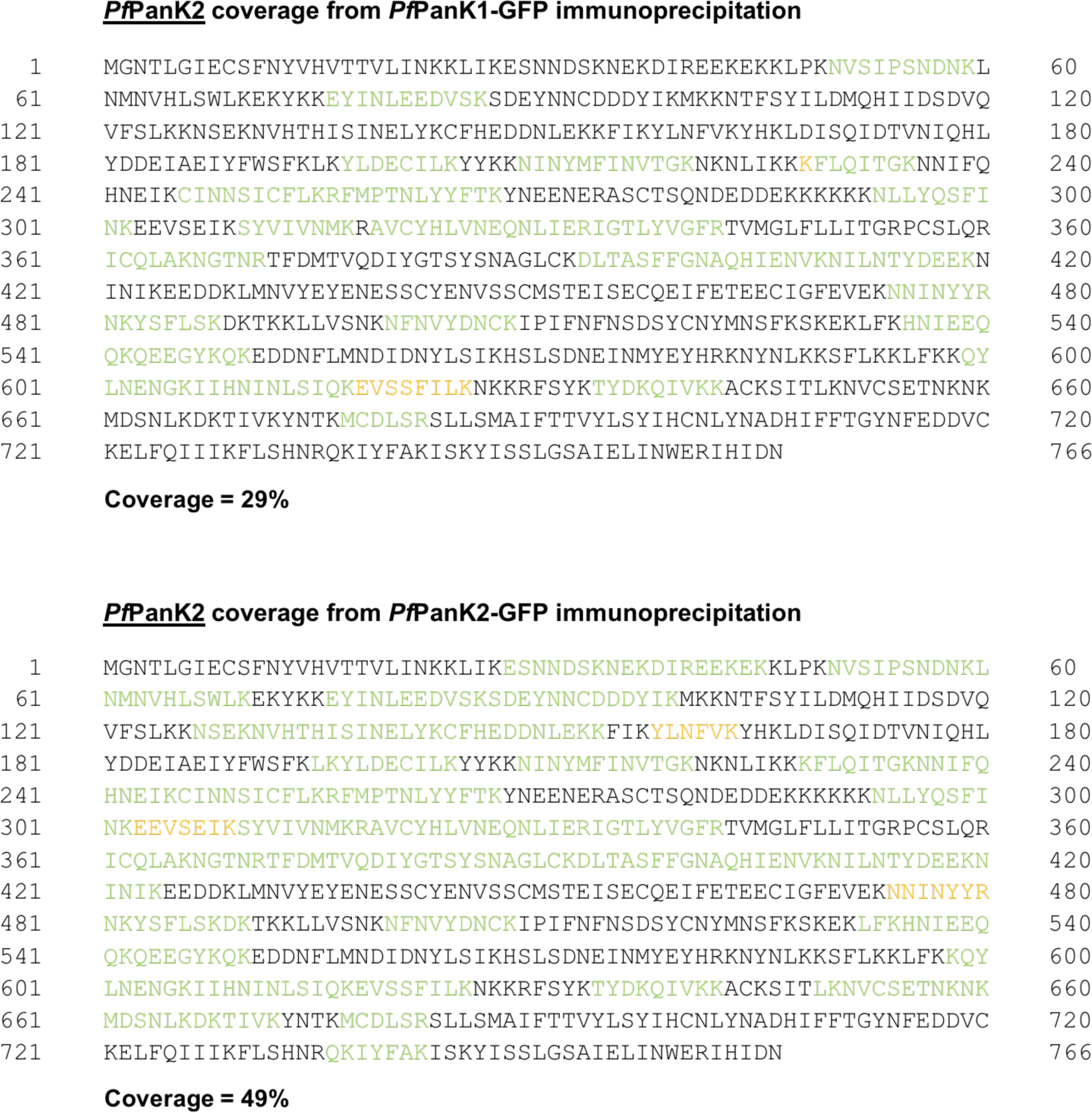
MS coverage of *Pf*PanK2. *Pf*PanK2 peptides detected in the two independent MS analyses of the GFP-Trap immunoprecipitation of the *Pf*PanK1-GFP- and *Pf*PanK2-GFP-expressing parasites. Residues in green were detected in either analysis with >95% confidence, while residues in orange were detected in either analysis with >90% (but <95%) confidence. Percentage coverage was calculated using only the residues labelled green.

**Figure S4.**
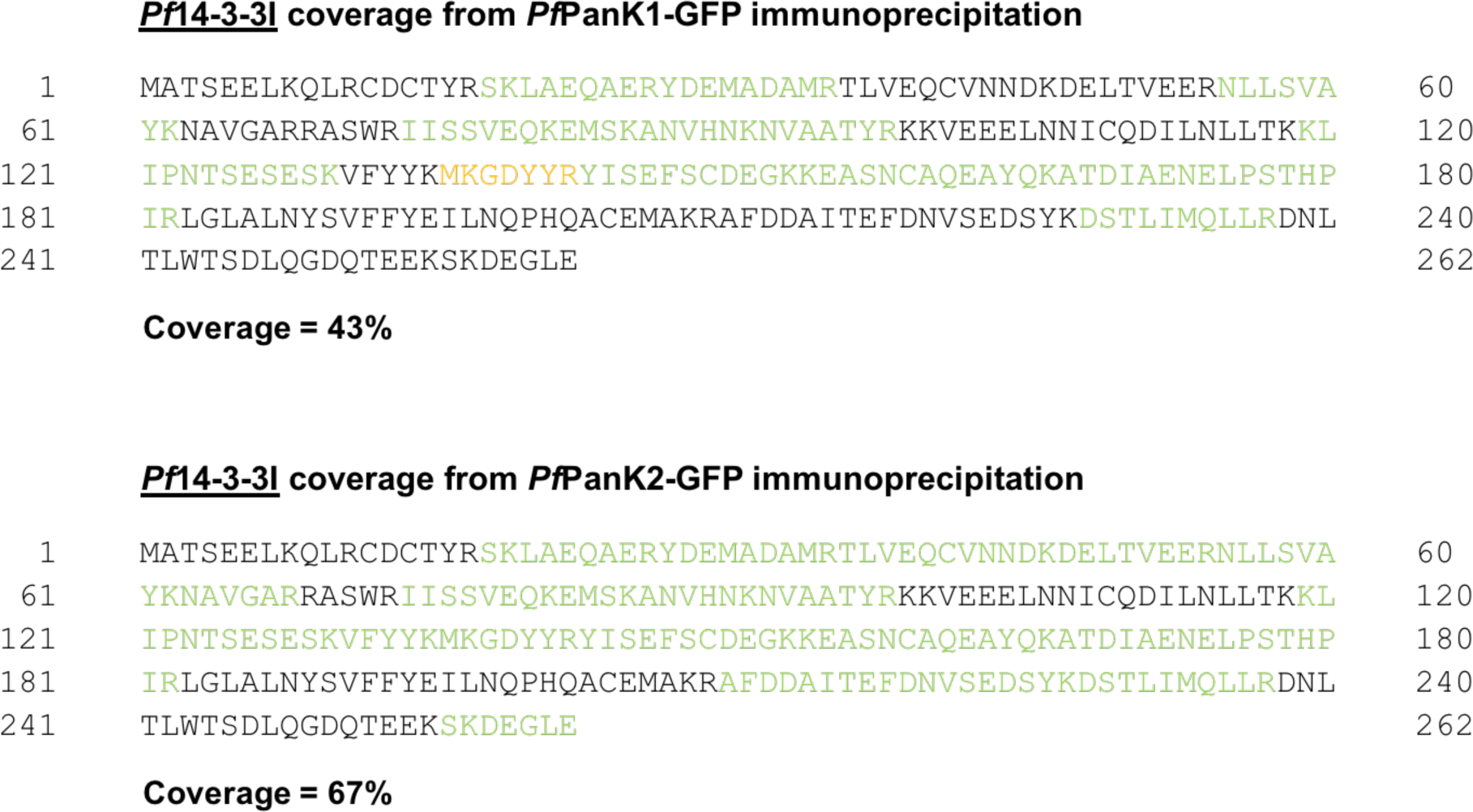
MS coverage of *Pf*14-3-3I. *Pf*14-3-3I peptides detected in the two independent MS analyses of the GFP-Trap immunoprecipitation of the *Pf*PanK1-GFP- and *Pf*PanK2-GFP-expressing parasites. Residues in green were detected in either analysis with >95% confidence, while residues in orange were detected in either analysis with >90% (but <95%) confidence. Percentage coverage was calculated using only the residues labelled green.

**Figure S5.**
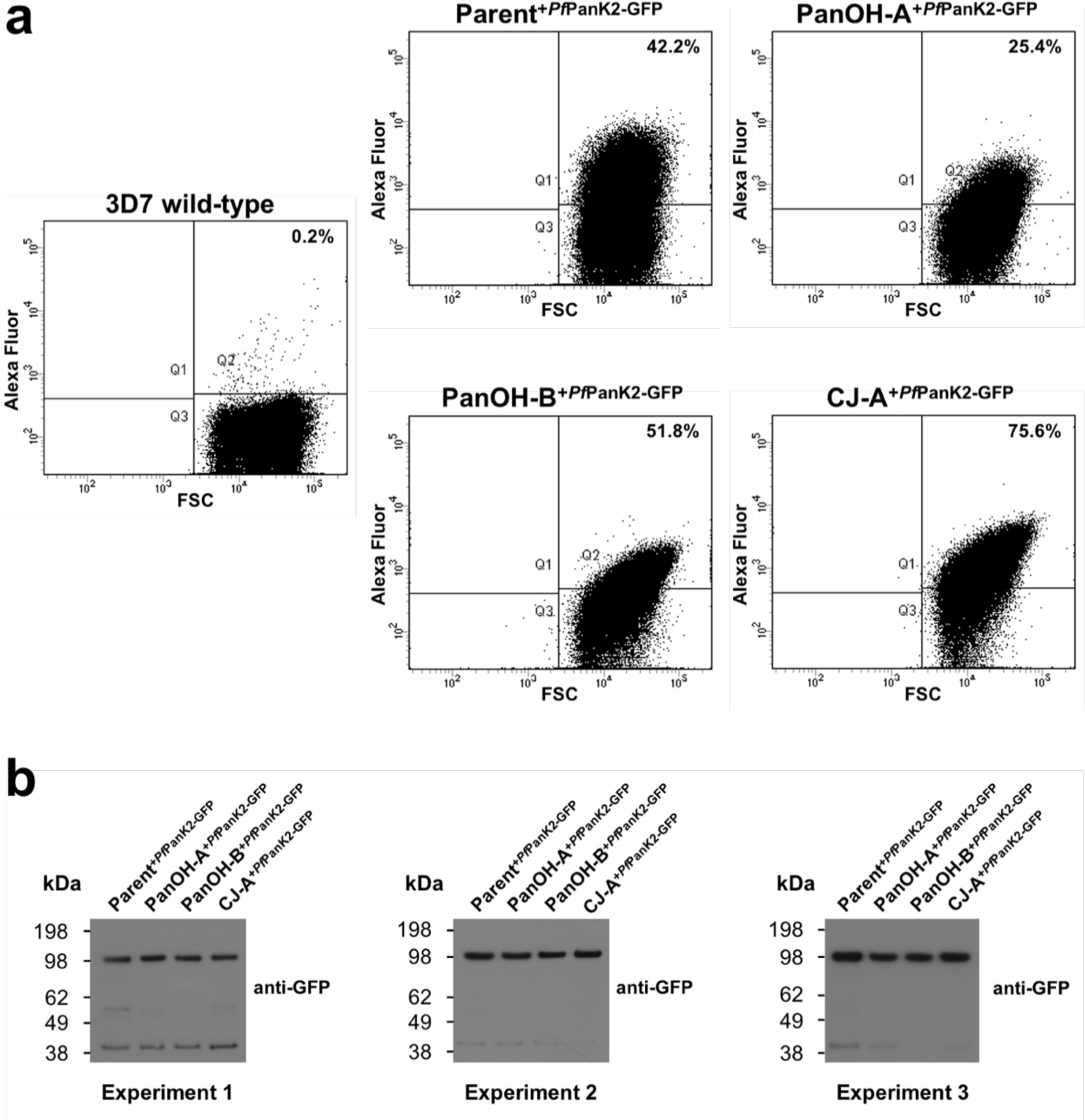
Determining the amount of GFP-Trap bound *Pf*PanK2-GFP for pantothenate phosphorylation assays. (a) The proportion of GFP-positive saponin-isolated 3D7, Parent^+*Pf*PanK2-GFP^, PanOH-A^+*Pf*PanK2-GFP^, PanOH-B^+*Pf*PanK2-GFP^ and CJ-A^+*Pf*PanK2-GFP^ trophozoites was determined by FACS analysis. The forward scatter (FSC) intensity on each x-axis corresponds to cell size and the AlexaFluor intensity on each y-axis corresponds to GFP fluorescence. The proportion of GFP-positive cells in each transgenic line (percentage value in each plot) was determined by using 3D7 trophozoites to set a gating threshold below which parasites were defined to be auto-fluorescent. Data shown are representative of three independent experiments, each performed prior to the [^14^C]pantothenate phosphorylation assays presented in Figure 2bii. The flow cytometry data was used to standardise the amount of *Pf*PanK2-GFP immunoprecipitated from each cell line used in each [^14^C]pantothenate phosphorylation assay. (b) Denaturing western blot analysis of *Pf*PanK2-GFP in the GFP-Trap immunoprecipitated complexes that were used in the [^14^C]pantothenate phosphorylation assays performed to generate the data in Figure 2b. Western blots were performed with an anti-GFP antibody and each blot shows the relative amounts of *Pf*PanK2-GFP immunopurified from the four different cell lines used in each of the three [^14^C]pantothenate phosphorylation experiment. The same volume of samples (10 µL per lane) was used for all three experiments.

**Figure S6.**
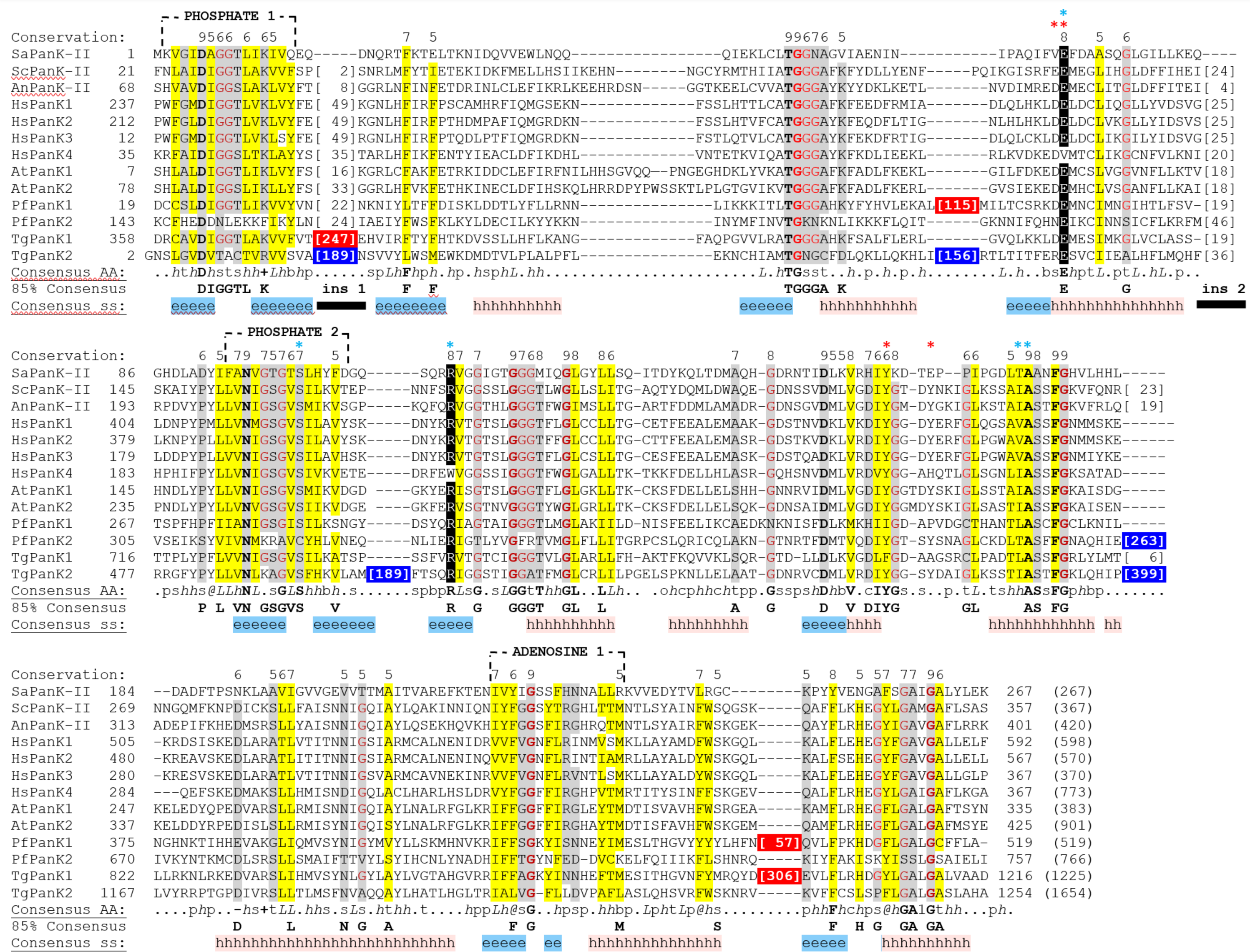
Multiple sequence alignment of representative Type II PanKs. The conserved PHOSPHATE 1, PHOSPHATE 2, and ADENOSINE 1 motifs of the acetate and sugar **kinases**/Hsc70/actin (ASKHA) superfamily of kinases are labelled at the top of the alignment. The Glu (E) residue involved in catalysis and the Arg (R) residue involved in positioning the substrate, are shown on a black background. Residues that have been found to interact with pantothenate and acetyl-CoA in human PanK3 ^9,10^ are marked with a blue asterisk. Residues that were found to interact to stabilise the human PanK3 active site are marked with a red asterisk. The catalytic Glu (E) residue is marked with a red and blue asterisk as it is involved in both the interaction with pantothenate and the stabilisation of the active site through interaction with a Tyr (Y) residue of the opposite protomer. The numbers at the start and end of each sequence indicate the position of the first and last residue in the alignment, respectively. The lengths of insertions are specified within the square brackets and the total length of protein sequences are shown in round brackets. Residues within the ASKHA superfamily motifs and conserved residues are highlighted based on the consensus AA guide for the column as follows: identical = bold, hydrophobic (W,F,Y,M,L,I,V,A,C,T,H) = yellow, charged/polar/small (D,E,K,R,H/D,E,H,K,N,Q,R,S,T/A,G,C,S,V,N,D,T,P) = grey and Gly = red. The two insertion regions (Ins 1 and Ins 2) common to eukaryotic type II PanKs, but absent in prokaryotic PanKs are indicated by the black horizontal bars, while the *Pf*Pank1/*Tg*PanK1 and *Pf*PanK2/*Tg*PanK2 specific inserts are highlighted on a red and blue background, respectively. Conservation refers to the conservation indices. Values at and above the conservation index cut-off (5) are displayed above the amino acid. Consensus AA: refers to the consensus level alignment parameters for the consensus amino acid sequence. This is displayed if the weighted frequency of a certain class of residues in a position is above 0.8. Consensus symbols: conserved amino acids are in bold and uppercase letters; aliphatic (I, V, L): *l*; aromatic (Y, H, W, F): *@*; hydrophobic (W, F, Y, M, L, I, V, A, C, T, H): *h*; alcohol (S, T): o; polar residues (D, E, H, K, N, Q, R, S, T): p; tiny (A, G, C, S): t; small (A, G, C, S, V, N, D, T, P): s; bulky residues (E, F, I, K, L, M, Q, R, W, Y): b; positively charged (K, R, H): **+**; negatively charged (D, E): **-**; charged (D, E, K, R, H): c. Marked below the alignment, 85% consensus includes those residues that occur in either the superfamily motifs and/or conserved residues where the same residue occurs more than 85% (10 out of 13 sequences). Consensus secondary structure (ss) elements: h = alpha helix, e = beta strand. Species names are abbreviated as follows: *Sa* = *Staphylococcus aureus*, *Sc* = *Saccharomyces cerevisiae*, *An* = *Aspergillus nidulans*, *Hs* = *Homo sapiens*, *At* = *Arabidopsis thaliana, Pf* = *Plasmodium falciparum* and *Tg* = *Toxoplasma gondii*. The alignment was created using PROMALS3D ^8^.

**Figure S7.**
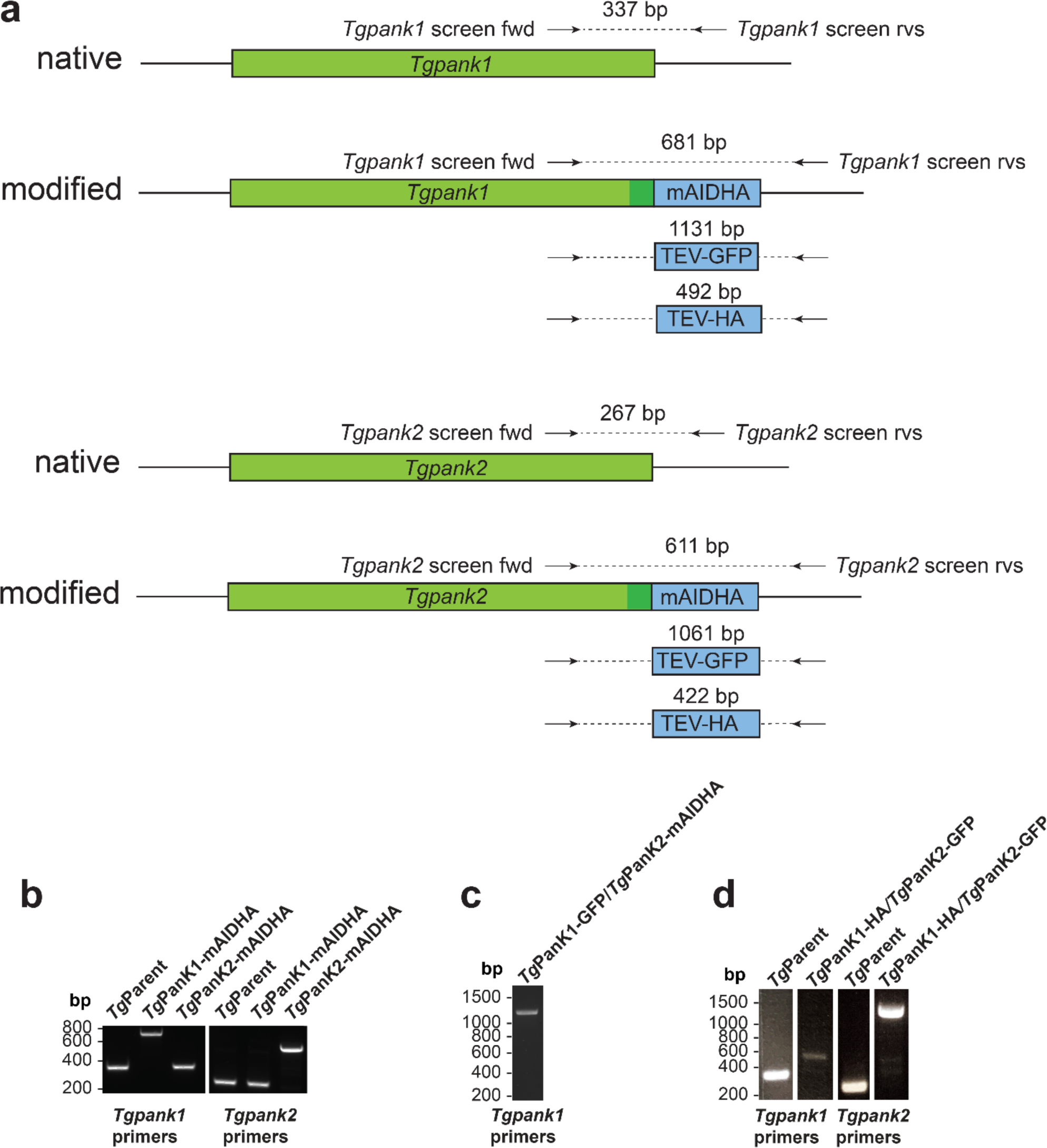
Gene models and confirmation of the incorporation of the coding sequence for various epitope tags into the *Tgpank1* and *Tgpank2* loci. (a) Gene models for *Tgpank1* and *Tgpank2* indicating the incorporation site of the epitope tag coding sequence. The expected sizes of the PCR products when screened with each set of screening primers are shown above the corresponding epitope tag coding sequence. The screening primers are *Tgpank1* screen fwd and rev for *Tgpank1* (referred to as *Tgpank1* primers in the panels b-d), and *Tgpank2* screen fwd and rev for *Tgpank2* (referred to as *Tgpank2* primers in panels b-d). Primers are detailed in Table S1. (b) PCR analysis of the *Tg*Parent, and singly-tagged *Tg*PanK1-mAIDHA and *Tg*PanK2-mAIDHA clonal lines. As can be seen, both *Tg*PanK1-mAIDHA and *Tg*PanK2-mAIDHA have successfully incorporated mAIDHA tags. (c) PCR analysis of the doubly-tagged *Tg*PanK1-GFP/*Tg*PanK2-mAIDHA clonal line. CRISPR/Cas9 was utilised to incorporate a sequence encoding a TEV-GFP tag into the genomic locus of the *Tgpank1* gene within the *Tg*PanK2-mAIDHA-expressing line. (d) PCR analysis of the *Tg*PanK1-HA/*Tg*PanK2-GFP doubly tagged clonal line (*Tg* Clone B4C6). CRISPR/Cas9 was utilised to incorporate a sequence encoding a TEV-HA tag into the genomic locus of the *Tgpank1* gene and a sequence encoding TEV-GFP tag into the genomic locus of the *Tgpank2* gene.

**Figure S8.**
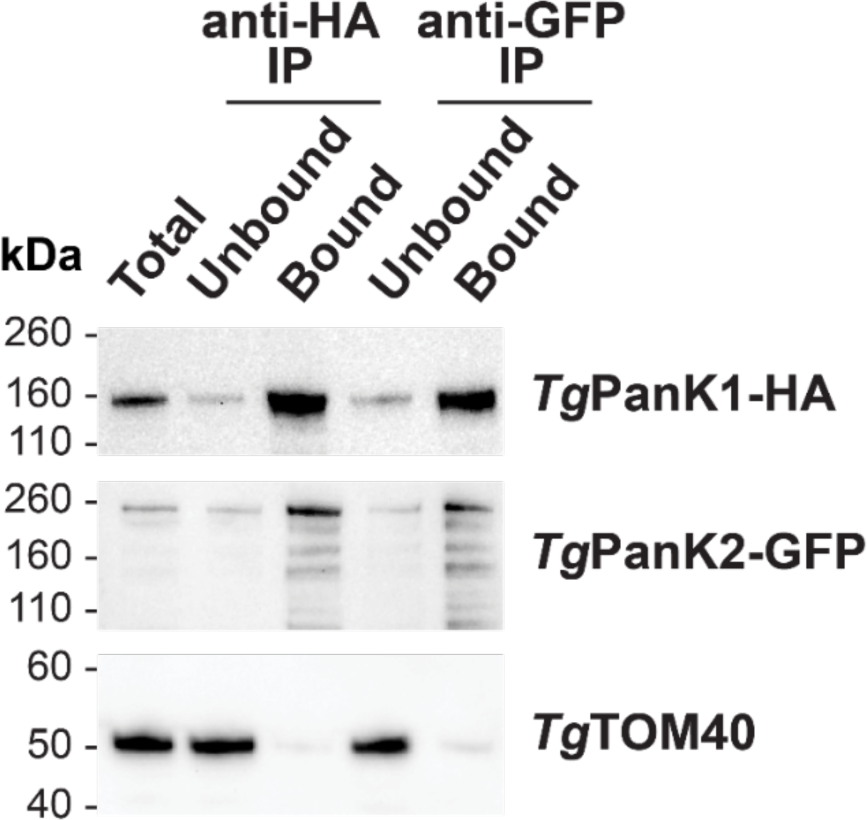
Anti-GFP and anti-HA immunoprecipitation of *Tg*PanK1-HA/*Tg*PanK2-GFP expressing parasites. Anti-HA and anti-GFP denaturing western blot analysis of fractions from GFP-Trap and anti-HA immunoprecipitations performed using lysates prepared from the parasite lines expressing *Tg*PanK1-HA/*Tg*PanK2-GFP. The expected molecular masses of *Tg*PanK1 and *Tg*PanK2 are ∼132 kDa and ∼178 kDa, respectively. The molecular mass of GFP is ∼27 kDa. The blot shown is representative of three independent experiments, each performed with different batches of parasites. Denaturing western blots were also probed with anti-*Tg*TOM40, which served as a loading control.

**Figure S9.**
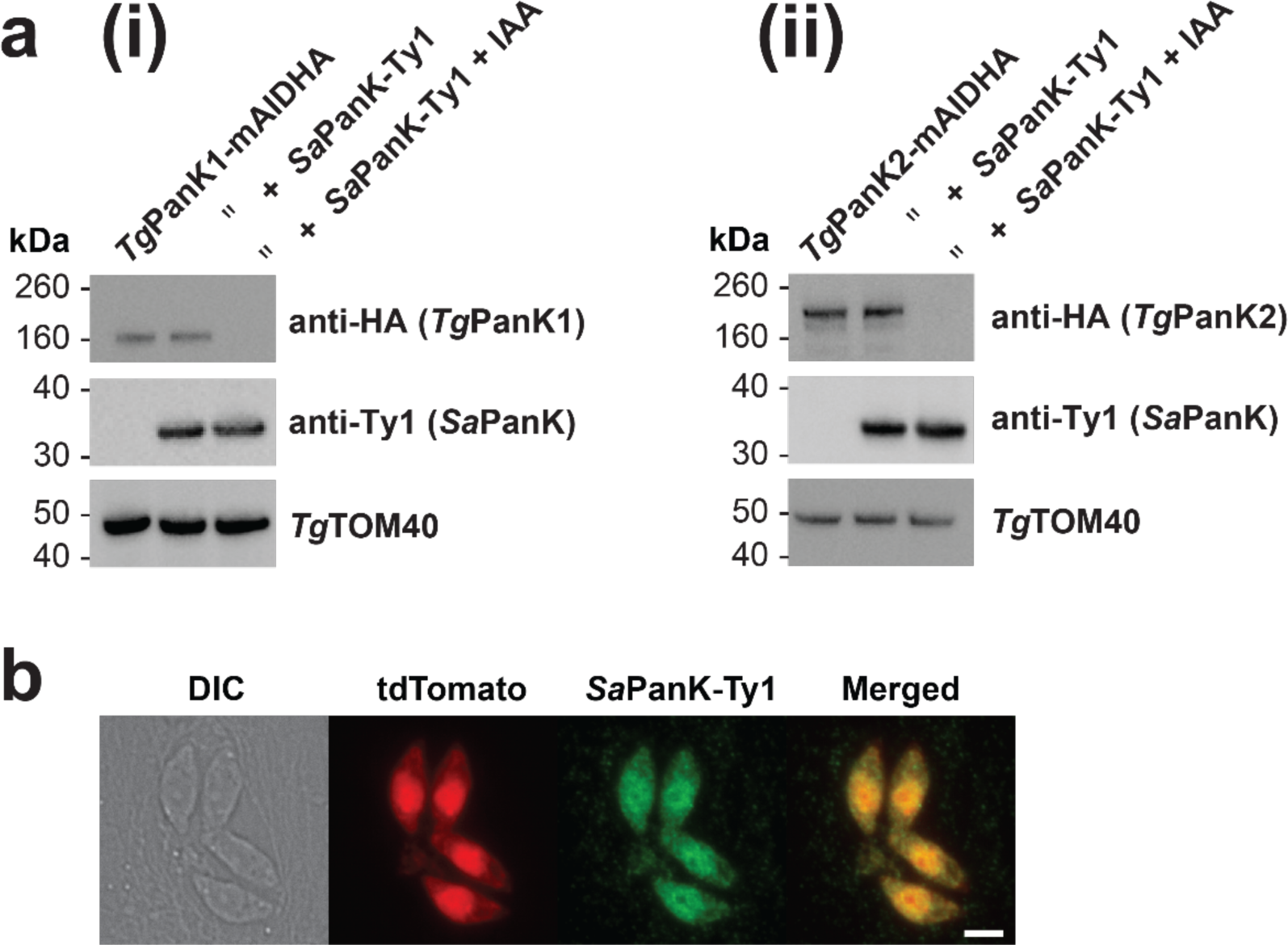
Expression of *Sa*PanK-Ty1 in *Tg*PanK1-mAIDHA- and *Tg*PanK2-mAIDHA-expressing parasites. (a) Anti-HA and anti-Ty1 denaturing western blot analysis of *Sa*PanK-Ty1-complemented and non-complemented (i) *Tg*PanK1-mAIDHA- and (ii) *Tg*PanK2-mAIDHA-expressing lines, in the absence or presence (for 1 h) of 100 µM IAA. The expected molecular masses of *Tg*PanK1-mAIDHA, *Tg*PanK2-mAIDHA and *Sa*PanK-Ty1 are ∼141 kDa, ∼187 kDa and ∼29 kDa, respectively. Denaturing western blots were also probed with anti-*Tg*TOM40, which served as a loading control. Each blot shown is representative of three independent experiments, each performed with a different batch of parasites. (b) Fluorescence micrographs of a HFF cell infected with four tachyzoite-stage *Tg*PanK1-mAIDHA^+*Sa*PanK-Ty1^ parasites within a vacuole, indicating the presence of *Sa*PanK-Ty1. From left to right: Differential interference contrast (DIC), tdTomato-fluorescence indicating the location of the parasites within the host cell, anti *Sa*PanK-Ty1 AlexaFluor 488 fluorescence, and merged images. Scale bar represents 2 µm.

**Figure S10.**
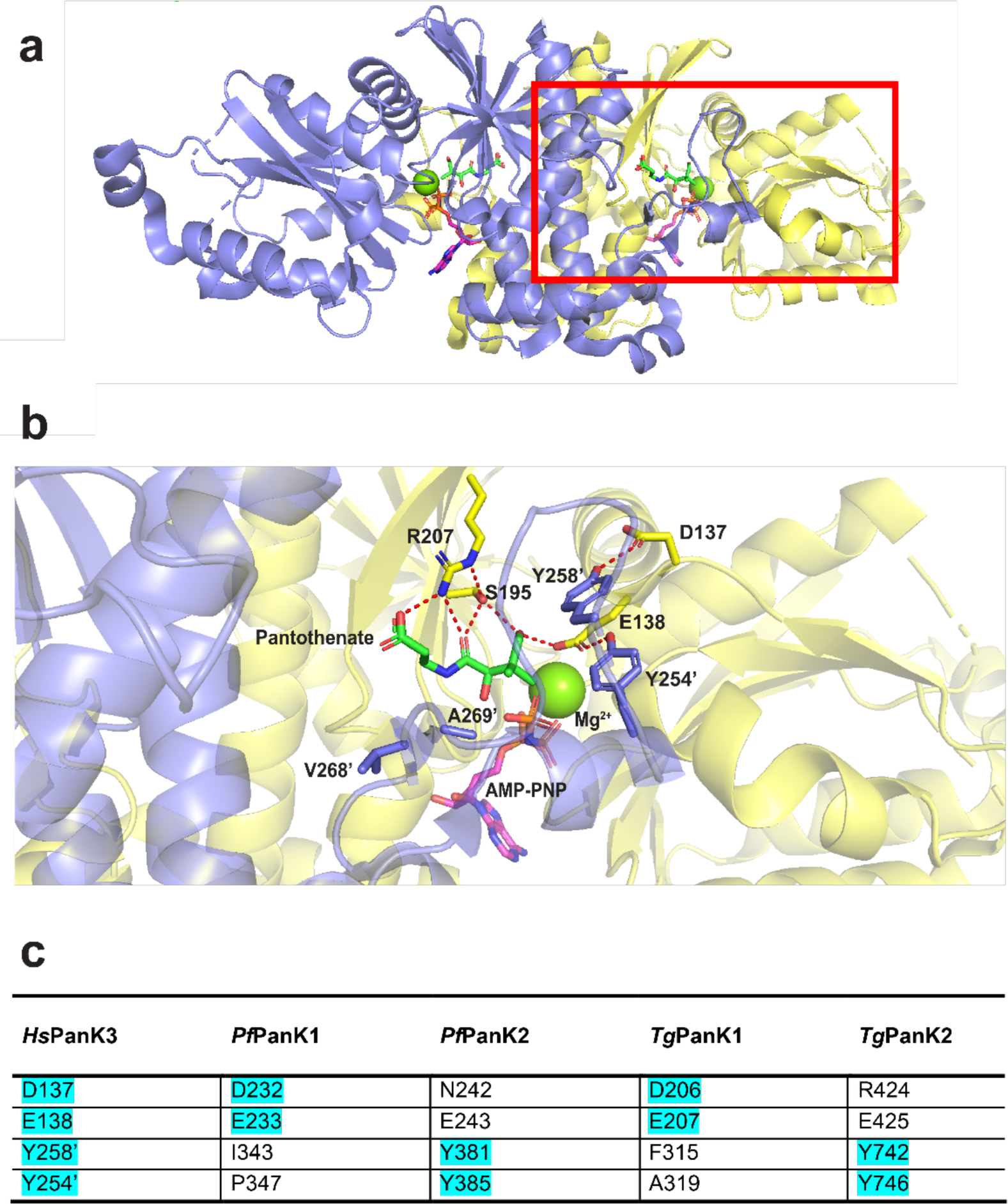
Pantothenate binding site and interactions in *H. sapiens* PanK3. (a) *H. sapiens* AMP-PNP-pantothenate-bound PanK3 crystal structure (PDB ID: 5KPR, Subramanian *et al.*^10^). The homodimeric protein is made up of two identical protomers (lilac and yellow) forming two identical active sites, each binding pantothenate (green). The red square encompasses one of the active sites. (b) Magnification of the region outlined by the red square in (a). Residues from both protomers contribute to the stabilisation of the binding pocket (E138 forms a hydrogen bond with Y254’ and D137 with Y258’) and interact with pantothenate (E138, S195, R207, A269’ and V268’). Hydrogen bonds with and between the sidechains of these residues are shown in red. An apostrophe denotes residues from the lilac protomer. (c) List of residues annotated in the *Hs*PanK3 model that participate in the stabilisation of the binding pocket (highlighted cyan), and a comparison to the equivalent residues in *P. falciparum* and *T. gondii* PanKs. The PanKs from *P. falciparum* and *T. gondii* do not individually contain the complete set of residues required for the stabilisation of the binding pocket, but the combination of residues (highlighted cyan) from PanK1 and PanK2 suggests that each PanK1/PanK2 heterodimer will have only one stabilised binding site.

**Figure S11.**
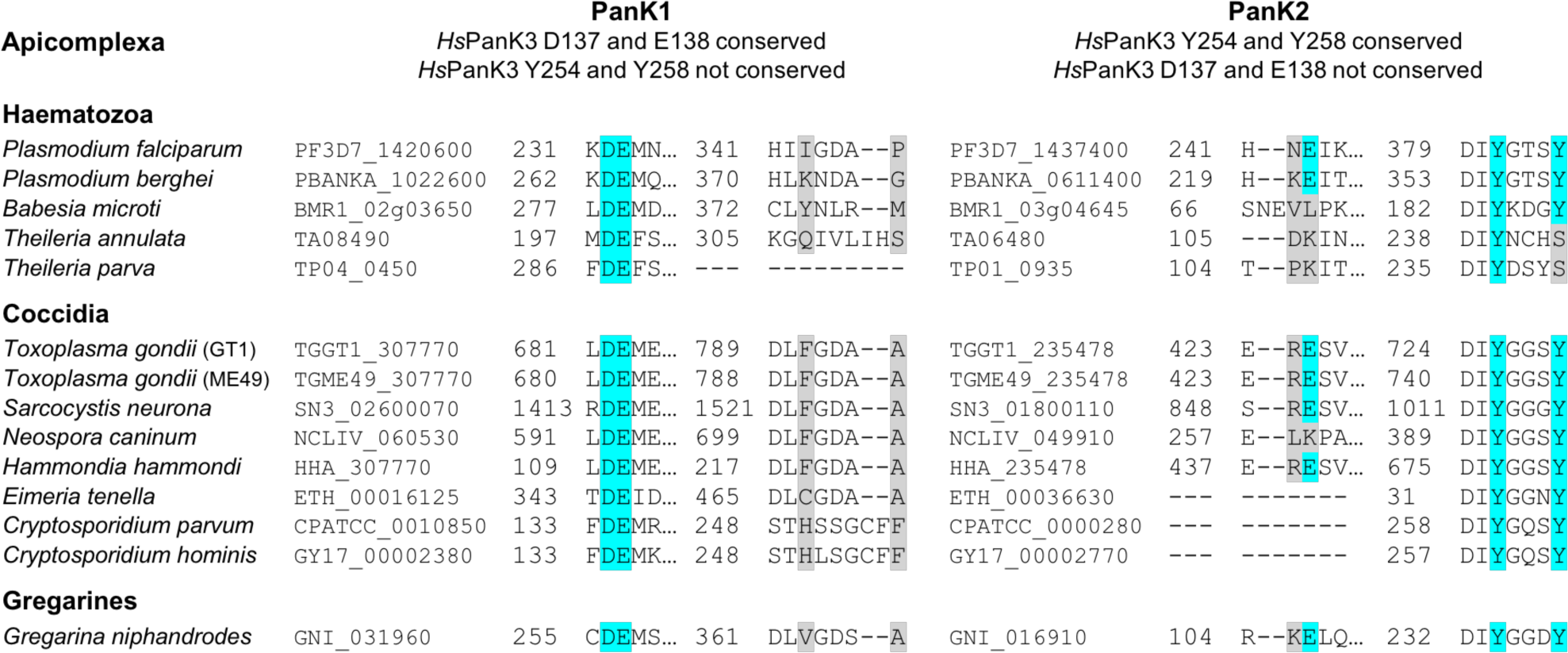
Multiple sequence alignment of active site stabilisation residues in apicomplexan PanKs. The apicomplexan PanK residues corresponding to the *Hs*PanK3 residues that are involved in the stabilisation of the binding pocket (D137, E138, Y254 and Y258) are highlighted in cyan if they are conserved and in grey if they are not conserved. The numbers before each alignment indicate the position of the first residue in the alignment. Each apicomplexan PanK is grouped into PanK1 or PanK2 based on their similarity to either *Pf*PanK1/*Tg*PanK1 or *Pf*PanK2/*Tg*PanK2, respectively. The alignment was created using PROMALS3D ^8^.

